# Filter-aided expansion proteomics

**DOI:** 10.1101/2023.11.29.569242

**Authors:** Zhen Dong, Yi Zhu, Fangfei Zhang, Chunlong Wu, Ting Chen, Jiayi Chen, Wenhao Jiang, Qi Xiao, Shu Zheng, Kiryl D. Piatkevich, Tiannan Guo

## Abstract

Integration of hydrogel-based tissue expansion and mass spectrometry-based proteomics enables high spatial resolution analysis of global protein expression in tissues. Here, we introduce a filter-aided tissue expansion proteomics (FAXP) method with optimized hydrogel preparation and treatment steps for spatial proteomics analysis of formalin-fixed paraffin-embedded (FFPE) specimens. Compared with our original ProteomEx method, FAXP, partially automated with robotics, exhibited 14.5-fold higher volumetric resolution, increased peptide yield leading to 250% more protein identifications, and 50% time consumption for sample preparation. We demonstrated its application in clinical colorectal tissue samples. We further integrated it with laser capture microdissection for proteomic analysis of subcellular organelles.

## Introduction

To study protein expression in space at the proteome level, two major strategies have been developed. The first one is mass spectrometry imaging (MSI)^1^, which usually use Matrix-Assisted Laser Desorption-Ionization (MALDI) to convert proteins or peptides in a particular region into gas-phase ions before MS analysis^2,3^. The other emerging spatial proteomics technologies are developed as the sensitivity of MS-based proteomics increases to single cell level. For example, selected regions of tissue could be isolated using laser-capture microdissection (LCM)^4–7^, which might be coupled with microfluidics^8,9^ or a 3D-printed device^10^ or tissue clearing^11^, followed by highly sensitive MS-based proteomics analysis. Proteins on the surface of selected tissue region could also be extracted using a fine liquid junction in line with LC-MS/MS hardware^12,13^.

Tissue expansion technology has been recently introduced to MS-based proteomics^14–17^. The concept of expansion proteomics (ProteomEx) is based on the expansion microscopy approaches^18–20^, like protein-retention Expansion Microscopy^21^, appended with MS-based proteomics analysis. First, paraformaldehyde (PFA)-fixed tissue section is treated with N-succinimidyl acrylate (NSA) to modify primary amino groups of proteins with acryl group, capable of co-polymerizing with hydrogel monomer. Then, treated tissue is infused with hydrogel monomers, and *in situ* polymerization reaction is initiated to form a tissue-hydrogel composite. To enable isotropic expansion, the formed tissue-hydrogel composite is mechanically homogenized by SDS-containing buffer at elevated temperatures (95°C). Expanded sample is manually microdissected, and excised pieces of hydrogel are subjected to in-gel reduction/alkylation and tryptic digestion to extract peptides for LC-MS/MS analysis. Importantly, chemical treatment of biological tissue does not introduce any modifications to the extracted peptides, yielding proteomics data indistinguishable from conventional sample preparation methods. Furthermore, unlike the LCM-based techniques, ProteomEx does not require any sophisticated or custom equipment and utilizes only commercially available and highly accessible reagents and accessories for sample preparation. However, the lateral and volumetric resolution of the original ProteomEx protocol remains limited to ∼160 µm and ∼0.61 nL, respectively^14^. Due to the extended sample treatment timeline (∼54 hours, not including MS analysis) and multiple manual steps, processing multiple samples in parallel is challenging, precluding large-scale analysis. Furthermore, the compatibility of ProteomEx with archived clinical samples was not previously demonstrated. Addressing these challenges and limitations of ProteomEx can significantly broaden its utility and promote large-scale spatial proteomic studies.

Here, we introduced a method termed filter-aided expansion proteomics (FAXP), compatible with multiple types of formalin-fixed paraffin-embedded (FFPE) tissues, including archived clinical samples. Compared to the original protocol, FAXP enables 2.2-/14.5-fold higher lateral/volumetric resolution with twice faster timeline. When integrated with laser capture microdissection, FAXP enables proteomics analysis of subcellular organelles. The FAXP method offers better reproducibility for subnanoliter sample volume and higher throughput, capable of processing 96 samples per batch across multiple batches in parallel. These enhancements stem from optimized procedures used in the sample expansion procedures and the development of filter-aided in-gel digestion for peptide retrieval. We further demonstrated its application in human colorectal cancer FFPE samples utilizing a hybrid library search strategy^22^ to identify 6790 proteins associated with the advancement of colorectal tumors through four stages alongside a machine learning classifier for feature selection and spatial visualizations of the specific features.

## Results

### Acceleration of the ProteomEx workflow

Since the ProteomEx method^14^ does not allow effective analysis of FFPE tissues, which are the most common specimens in clinic, we sought to optimize and enhance the original ProteomEx protocol to achieve several key features, including 1) compatibility with archived clinical samples; 2) faster timeline; 3) higher sample preparation throughput; 4) higher lateral and volumetric resolution. To mimic the popular format of archived clinical samples, we used 10 µm FFPE mouse liver tissue (unless otherwise stated), since it is one of the most homogenous tissues in terms of protein distribution, providing less biologically biased comparison between small samples across tissue sections^5,23^.

First, we asked whether the timeline of sample preparation and peptide recovery can be accelerated without sacrificing proteome coverage. We found that protein anchoring, hydrogel embedding, homogenization, and Coomassie staining, the most time-consuming steps in the original protocol, could be significantly accelerated (in total to 11.2 hours vs. 40.5 hours in ProteomEx) without compromising peptide and protein identifications compared to ProteomEx (Figure S1; see Methods section for the protocol details). However, we noticed that autoclavation of tissue-hydrogel composite at 105-120°C, which we employed for rapid homogenization, resulted in softer hydrogels after full expansion compared to the original homogenization treatment conditions. As softer hydrogels were harder to handle during dissection step and could potentially introduce larger distortion to the expanded tissue, we increased the hydrogel cross-linker ratio by 17.5 folds (Table S1), which enhanced hydrogel sturdiness similarly to that observed for other hydrogels reported before^24^. To evaluate tissue expansion using the altered hydrogel composition, we quantified the linear expansion factor and isotropy of expansion on cellular and whole tissue slice level using protein anchoring time as a variable (Figure 1A-G). At the cellular level, under 1-hour anchoring condition the linear expansion factors of cell nuclei visualized via DAPI staining were comparable to those measured at the whole expanded tissue (linear expansion factor (LEF) 4.42±0.31 vs. 4.69±0.17 for individual nuclei vs. whole expanded tissue, respectively; Figure 1E), indicating a consistent and uniform expansion pattern that extends from the cell nuclei to the whole tissue structure. However, extended anchoring time (2 hours) resulted in reduced expansion factors (LEF=3.10±0.58, p=4.50e-6) at the cellular level compared to those measured at the whole expanded tissue level (Figure 1E). Using 1-hour anchoring step for all further experiments, we computed the root-mean-square (RMS) measurement error in feature-length measurements after non-rigid registration of pre- and post-expansion tissue across length scales of up to 1500 µm. The analysis revealed that the RMS errors were approximately 5.5% of the measurement distance, which was slightly better than that for the original protocol (Figure 1F, G). The comparable RMS errors were also measured for tissues with rich extracellular matrix such as kidney (Figure S2), indicating a high degree of accuracy suitable for precise mapping of spatial proteome distribution onto pre-expanded tissue morphology.

**Figure 1.**
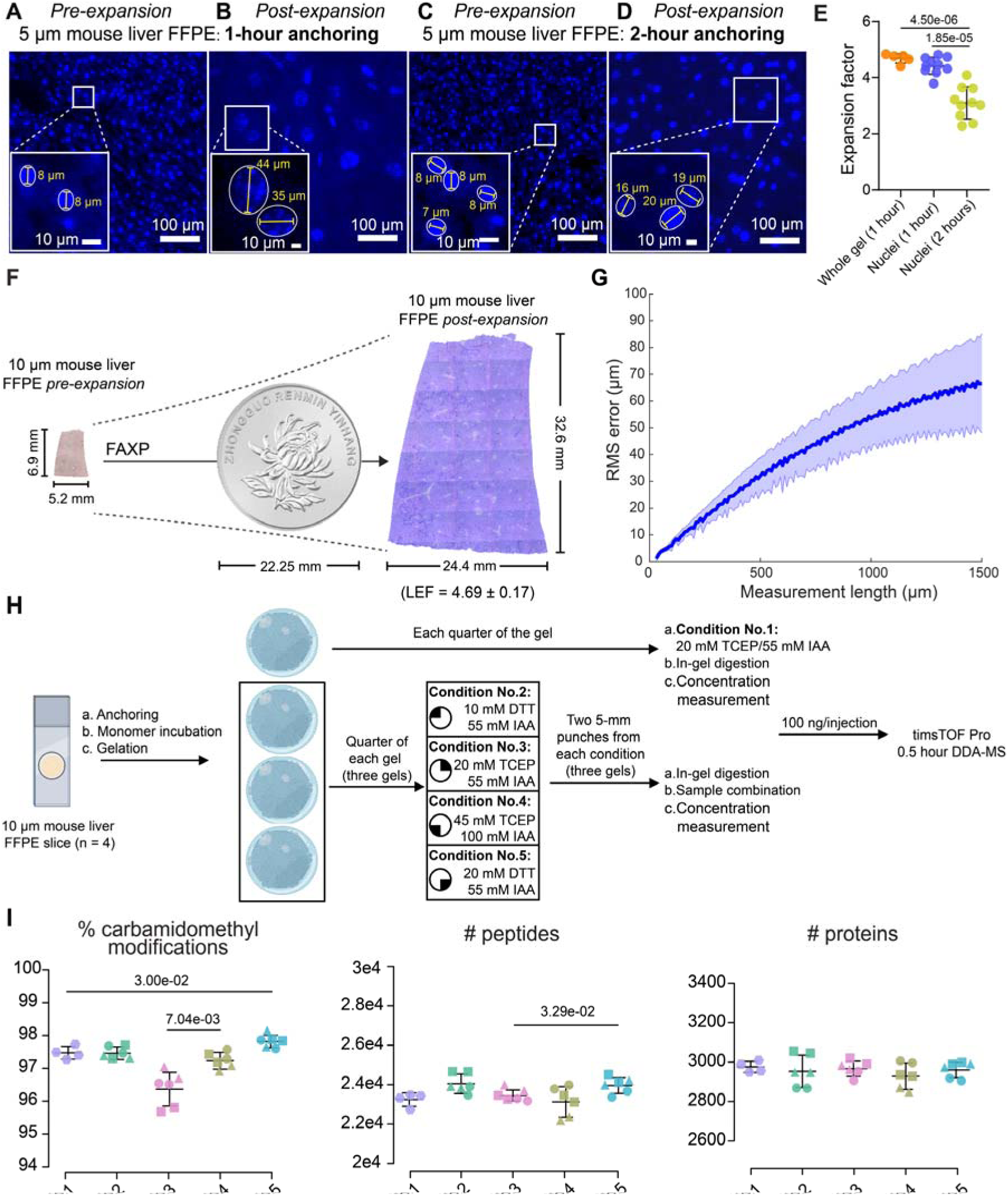
Optimization of gel-making procedures. (A) DAPI image of a mouse liver FFPE slice after 1-hour anchoring before expansion. (B) Post-expansion DAPI image of the same sample incubated under 1-hour anchoring. (C) DAPI image of a mouse liver FFPE slice after 2-hour anchoring before expansion. (D) Post-expansion DAPI image of the same sample incubated under 2-hour anchoring. Magnified views with white-circled nuclei and measured diameters are shown within each figure (A-D). Scale bars are indicated within each figure. (E) Comparison of expansion factors between whole gel macro-level colorimetric staining (n = 5) and micro-level DAPI nuclear staining under 1- and 2-hour anchoring (n = 10 each). All data points are presented as mean values ± standard deviation (SD). (F) Representative bright-field images of an FFPE mouse liver slice captured before expansion (left) and after expansion with Coomassie brilliant blue staining (right), in comparison to a one-dollar RMB coin, with a diameter of 22.25 mm. (G) Comparison of root-mean-square (RMS) measurement length error between pre-expansion and post-expansion liver slice images (n = 4); one example is shown in (F). (H) Study design for comparative proteomic analysis of four reduction and alkylation conditions. After gel-making for three mouse liver FFPE slices, four reduction/alkylation conditions including 10 mM DTT/55 mM IAA, 20 mM DTT/55 mM IAA, 20 mM TCEP/55 mM IAA, and 45 mM TCEP/100 mM IAA are applied to one-fourth of each gel (n = 3). Following in-gel digestion, samples from two 5-mm punches are mixed, and MS data is collected using 100 ng per injection. (I) Comparative proteomic results of the four reduction and alkylation conditions, including percentages of carbamidomethyl modifications, as well as the number of identified peptides and proteins. All data points are presented as mean values ± SD.

We then accelerated the ProteomEx timeline and streamlined peptide recovery from microdissected tissue-hydrogel samples, addressing the most laborious and unreliable steps due to multiple manual procedures (see Note S1, Figure S3, Figure S4, and Table S2 for details). 20 mM DTT/55 mM IAA can now be performed on the entire tissue section after homogenization, reducing the need for extra manual treatment steps during in-gel digestion and streamlining the protocol.

### Comparison of conventional in-gel digestion with filter-aided in-gel digestion

The original ProteomEx protocol had a limited lateral/volumetric resolution, achieving only ∼160 µm/∼0.61 nL, primarily due to difficulties in manually handling hydrogel samples smaller than 1 mm. Specifically, during peptide recovery small hydrogel pieces (<1 mm in dimension) underwent shrinkage caused by multiple buffer exchange steps resulting in attendant sample lost during pipetting. To overcome this limitation, we suggested immobilizing the hydrogel sample on a membrane placed into a pipette tip. We first assessed the feasibility of the proposed filter-aided in-gel digestion method (also referred to as in-tip digestion) compared to the original in-gel digestion (Figure 2A). Analysis of 1.55 nL of the tissue using timsTOF Pro 1-hour DDA-MS revealed that the in-tip digestion provided significantly higher number of peptide and protein identifications compared to in-gel digestion (increased by 97.7% and 30.6%, respectively; Figure 2B). Correspondingly, 75.0% of peptides and 97.5% of proteins identified in in-gel digestion were also identified in in-tip digestion, and it offered an extra identification of 56.7% more peptides and 20.5% more proteins, indicating a high consistency and performance improvement in proteome coverage (Figure 2C). In-tip digestion also displayed superior capability in identification of longer peptides (Figure 2D) without preference for peptide hydrophobicity (Figure 2E), indicating an improved peptide coverage for in-tip digestion compared with in-gel digestion. Furthermore, the in-tip digestion led to a significant improvement in protein digestion efficiency measured by missed cleavage rate, with an average of around 17.2%, compared to the in-gel digestion rate of 23.8% (Figure 2F). As an additional advantage, in-tip digestion successfully identified more mouse liver-specific markers^25^, including all discovered through in-gel digestion, and additionally six markers (ELOVL6, SLC35B1, PVRL1, TM6SF2, ABCA8A, and ADIPOR2) with lower abundance (Figure 2G). In-tip digestion generated more reproducible quantification outcomes with a significant improvement in coefficient of variance (less than 0.2 at peptide and protein levels; Figure 2H) and Pearson correlation (close to 1.0 at peptide and protein levels; Figure 2I). Both protocols exhibited analogous distributions of protein types and subcellular localizations as revealed by the pathway enrichment analysis using the identified proteins, indicating there was no significant biological variations from in-tip digestion (Figure 2J). Altogether, these results confirmed that filter-aided in-gel digestion method improved sample manipulation while enhancing coverage and reproducibility of proteome analysis.

**Figure 2.**
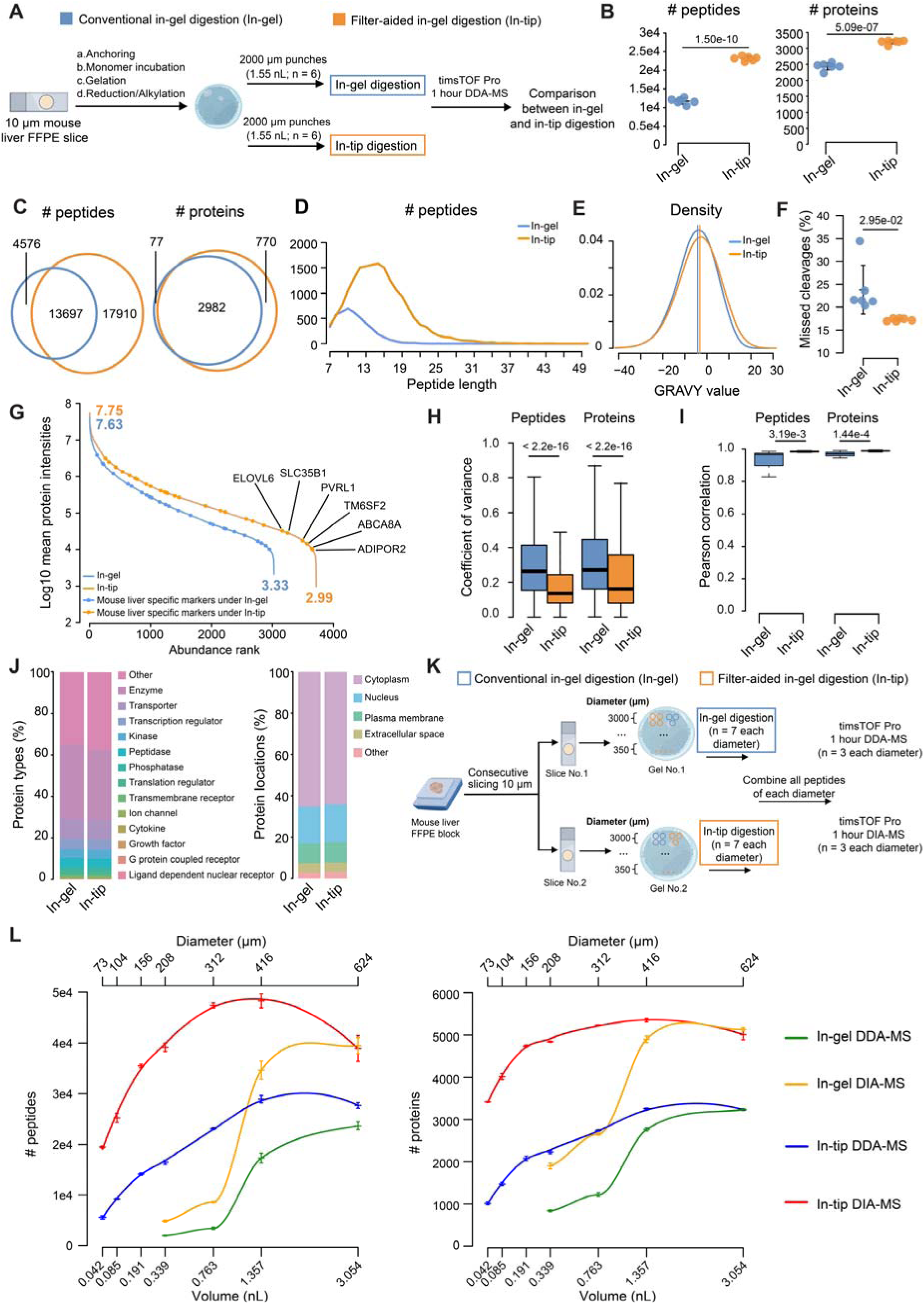
Comparison between conventional in-gel digestion and filter-aided in-gel digestion applied to mouse liver FFPE slices. (A) The study design illustrates a comparative proteomic analysis between conventional in-gel digestion (in-gel digestion) and filter-aided in-gel digestion (in-tip digestion). After gel-making for a mouse liver FFPE slice, 2000-µm punches (n = 6; containing 1.55 nL of tissue volume before expansion) are subjected to in-gel and in-tip digestions separately. (B) Number comparison of the identified peptides and proteins between in-gel and in-tip digestions (n = 6 each). All data points are presented as mean values ± standard deviation (SD). (C) Venn diagram shows the overlapping of identified peptides and proteins between in-gel and in-tip digestions. (D) A curve plot shows the distribution of peptide lengths exclusively identified through in-gel and in-tip digestions, as shown in (C). (E) A curve plot displays the density distribution of hydrophobicity (GRAVY) for identified peptides between in-gel and in-tip digestions. (F) Missed cleavage rates of the identified peptides using in-gel and in-tip digestions. (G) A curve plot shows log10-transformed mean protein intensities ranked by protein abundance. Mouse hepatocyte-specific markers are shown as circles. Markers exclusively identified under in-tip digestion are specifically labeled. (H) Coefficient of variation of the quantified peptides and proteins abundance from in-gel and in-tip digestions. (I) Pearson correlation for peptide and protein quantification from in-gel and in-tip digestions. (J) The protein types and subcellular locations of the identified proteins enriched through Ingenuity Pathway Analysis (IPA) are presented. (K) Study design presents a comparative proteomic analysis among different tissue volumes processed by in-gel and in-tip digestions. Gels are prepared for two consecutive mouse liver FFPE slices, and punches (n = 7) for each of the diameters (350 µm, 500 µm, 750 µm, 1000 µm, 1500 µm, 2000 µm, and 3000 µm) are collected separately for in-gel (1000 µm - 3000 µm) and in-tip (350 µm - 3000 µm) digestions. The combined peptides prepared under each diameter using different methods are acquired with 1-hour data-dependent acquisition mass spectrometry (DDA-MS; n = 3) and data-independent acquisition mass spectrometry (DIA-MS; n = 3), respectively. (L) The number of identified peptides and proteins for different tissue volumes with corresponding diameters shown in (K). The samples are processed by in-gel and in-tip digestions, and MS data are collected by DDA-MS and DIA-MS methods, respectively. All data points are presented as mean values ± SD.

### Improving volume-dependent limit of tissue micro-sampling

The combination of filter-aided in-tip digestion with accelerated ProteomEx workflow enabled improved proteome coverage while increasing the throughput of sample preparation. To validate whether in-tip digestion improves lateral/volumetric resolution of ProteomEx, we investigated the volume-dependent limit of tissue microsampling using the optimized workflow with peptide recovery step performed following either in-gel or in-tip protocol (Figure 2K). By manually dissecting tissue microsamples with actual volumes ranging from 0.042 nL to 3.054 nL (corresponds to lateral resolution from 73 µm to 624 µm), we observed that in-tip digestion under DDA-MS and DIA-MS identified a range of 5519 to 28,931 peptides and 19,499 to 48,330 peptides, respectively (Figure 2L). In-tip digestion consistently exhibited superior peptide and protein identification performance compared to in-gel digestion in both DDA and DIA modes, except for when processing tissue samples with a volume of 3.054 nL, indicating in-tip digestion was more prone for minute sample preparation. There was a saturating plateau observed for tissue volumes between 0.763 nL and 1.357 nL using in-tip digestion with a leveling off at higher volumes. In turn, due to sample loss during peptide extraction in-gel digestion encountered a technical constraint of processing tissue volumes below 0.339 nL, which was 8 times larger than the smallest volume we were able to process with in-tip digestion. For tissue volume of 0.339 nL (the smallest volume we could process with in-gel digestion), in-tip digestion outperformed in-gel digestion by around 250% in protein identification resulting in identification of 2229 and 4844 proteins in DDA and DIA mode, respectively. Remarkably, in-tip digestion displayed compatibility with tissue samples as small as 0.042 nL in volume, effectively achieving a lateral resolution of 73 µm on 10 µm thick sections, which was equivalent to 8 murine hepatocytes^26^. These results demonstrated that in-tip digestion improved lateral and volumetric resolution by 2.85 and 8.07-fold, respectively, while providing 8.07/2.55-fold higher peptide/protein identification compared to the ProteomEx protocol with in-gel digestion under its lower limit tissue volume using DIA mode.

In addition, the approach effectively targets single nucleus by combining LCM (Figure 3 and Supplementary Video 1), resulting in detection of an average of 550 proteins with an Astral mass spectrometer. When compared to the 0.042 nL In-tip DIA-MS (Figure S5), this method identified 258 specific proteins (Figure S5A) and demonstrated significantly higher ratios for nucleus and transcription regulator-related proteins, exhibiting a 4.26- and 3.59-fold increase, respectively (Figure S5B). Furthermore, the nuclei-specific pathway enrichment analysis revealed elevated enrichment in nucleic activities (Figure S5C), such as mRNA processing and splicing pathways.

**Figure 3.**
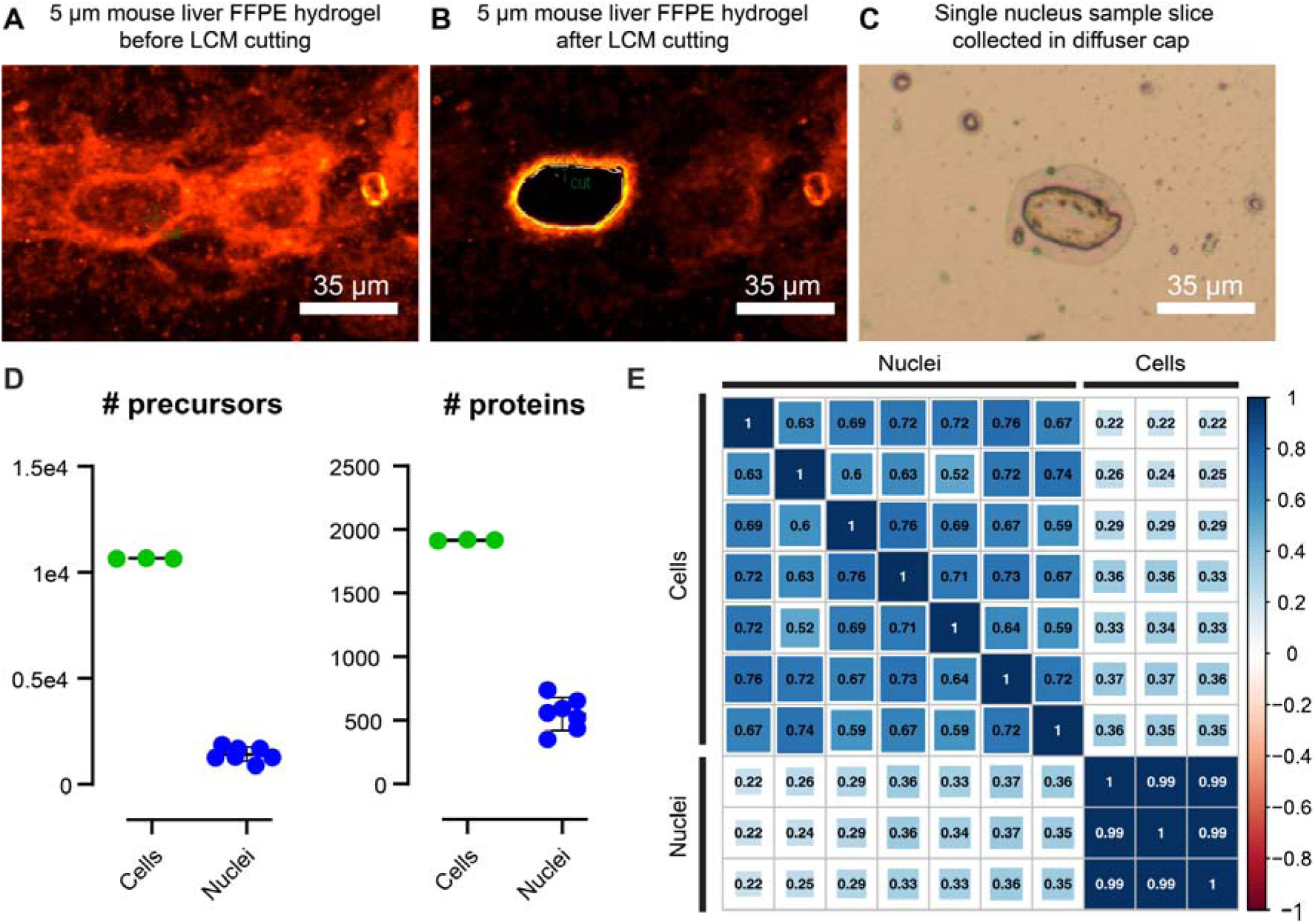
Study of single nucleus collection and analysis through tissue expansion, laser capture microdissection and filter-aided in-gel digestion. (A) SYPRO Ruby image of a mouse liver FFPE slice after expansion before laser capture microdissection (LCM) single nucleus cutting. (B) SYPRO Ruby image of the same sample after LCM single nucleus cutting. (C) Bright field image of the single nucleus in the diffuser cap. Scale bars are indicated within each figure. (D) Number of peptide and protein identifications between mouse hepatocytes and single nucleus in DIA mode through library-free search (data for mouse hepatocytes is from 0.042 nL of In-tip DIA-MS in Figure 2L, n=3; data for single nucleus is n=7 biological replicates from one liver slice). Data are presented as mean values ± SD. (E) Heatmap of Pearson correlations for protein quantification for each sample pair from mouse hepatocytes and nuclei. The color bar indicates the values of Pearson correlations.

### Application of FAXP on colorectal cancer patient samples

To further explore the utility of FAXP on clinical samples, we examined the proteome profiles of various subtype regions in colorectal cancer (CRC), namely N (normal), L (low-grade dysplasia), H (high-grade dysplasia), PC (peripheral carcinoma), CC (central carcinoma), and C (carcinoma), delineated using H&E staining by expert pathologist, aiming to examine spatial discrepancies at an individual patient level and evaluate variations within the same and between different subtypes across patients. We first confirmed the FAXP protocol enabled efficient Coomassie staining and isotropic expansion of the archive clinical samples (Figure 4A). Notably, the RMS error for CRC tissue was approximately 6% of the measurement distance, which was within the expected range based on mouse tissue expansion (Figure 4B). Subsequently, we performed visual-guided manual microdissection of the expanded CRC samples from three distinct patients (referred to as P1, P2, and P3) and collected samples as outlined in Figure 4C.

**Figure 4.**
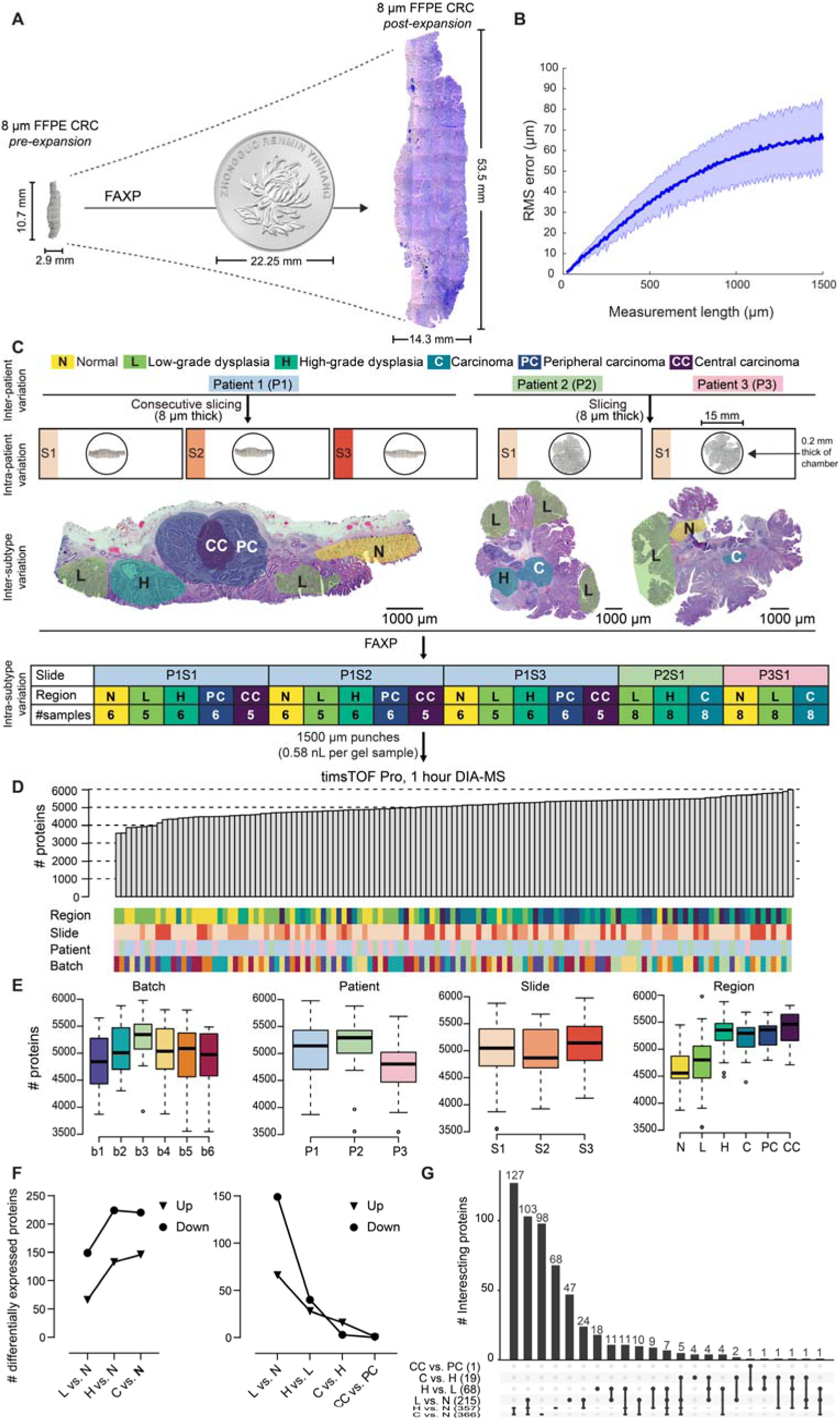
Application of FAXP on colorectal cancer patient samples. (A) Representative bright-field images of human colorectal cancer (CRC) slices before (left) and after (right) expansion using FAXP, in comparison to a one-dollar RMB coin, with a diameter of 22.25 mm. Expanded slices are stained with Coomassie brilliant blue for visualization. (B) Quantitative comparison of root-mean-square (RMS) measurement length error between original CRC slice images and corresponding expanded images (n = 3); one example is shown in (A). (C) Study design for the CRC application. Four different variations of patient samples are included - inter-patient variation: analysis of samples from three different patients (P1, P2, P3) to assess protein expression differences across individuals; intra-patient variation: examination of longitudinal variations using three consecutive slides from the first patient (P1S1, P1S2, P1S3); inter-subtype variation: investigation of proteomic changes across disease progression subtypes (N - normal, L - low-grade dysplasia, H - high grade dysplasia, C - carcinoma, PC - peripheral carcinoma, CC - central carcinoma); intra-subtype variation: exploration of protein expression variations within a specific disease subtype. H&E staining is performed on FFPE sample slides from three patients (P1, P2, P3). The slice IDs (P1S1, P1S2, P1S3, P2S1, P3S1) are associated with specific subtypes. The number of mass spectrometry (MS) samples for each subtype associated with each slice are indicated. 1500- µm punches (containing 0.58 nL of tissue volume before expansion) are subjected to filter-aided in-gel digestion and MS data are collected by timsTOF Pro using 1 hour DIA-MS. (D and E) Distribution of identified proteins are presented based on MS batches, patient numbers, slide numbers, and subtype regions, respectively. (F) Number of differentially expressed proteins (DEPs) in different comparisons. (G) Upset diagram showing the DEPs overlaps for selected comparisons.

In total 131 microdissected tissue samples were processed using the FAXP protocol and analyzed with optimized diaPASEF-based MS settings (Table S3). Considering the protein identification, we opted for the hybrid-pan human hybrid library for subsequent quantitative analysis due to its capability to generate the highest number of identifications, resulting in the recognition of 58,038 precursors and 6790 proteins (Figure S6E). To probe differences among various groups, we categorized the identified proteins based on MS batch IDs, patient IDs, slice IDs, and disease subgroups (Figure 4D). Our results revealed a consistent number of protein identifications across batches, patients, and slices (Figure 4E; Figure S6F-G), demonstrating the reproducibility and robustness of the FAXP workflow. The systematic data analysis revealed that the number of identified proteins progressively increased with the severity of cancer stage. Specifically, number of protein identifications in L, H, C, PC, CC were 2.50%, 14.15%, 12.23%, 14.62%, and 16.61% higher, respectively, than that in normal tissue (p = 0.36, 8.76e-08, 1.82e-05, 7.79e-09 and 1.67e-07), with a trend for increasing number of proteins with cancer progression (Figure 4E).

In the context of quantifying differences between various stages in CRC, the study identified significantly differentially expressed proteins (DEPs) when comparing both progression stages to normal stage and adjacent stages in the progression process (Figure 4F-G; Figure S6H). Notably, there were more down-regulated DEPs observed in more advanced stages compared to less severe stages, although this trend did not hold for comparisons such as C vs. H and CC vs. PC (Figure 4F). The number of DEPs found aligned with the increasing identification difference observed during the progression stages (Figure 4E). Specifically, when examining more severe stages such as C and H compared to the normal stage, there was a similarity in the most regulated DEPs. These included upregulated DEPs such as CEAM5, IFM3, and S100P, as well as downregulated DEPs such as CLCA1 and FCGBP. In the context of tumor development regulation, CEAM5 was noted as an oncogene that promotes tumor progression and induces resistance to anoikis in colorectal carcinoma cells^27^. The expression of CEAM5 exhibited an increasing trend in pair-wise comparisons such as L vs. N (FC = 2.43, p = 1.84e-03), H vs. N (FC = 10.17, p = 3.13e-09), and C vs. N (FC = 12.26, p = 3.13e-09). Besides, CLCA1, recognized as a tumor suppressor in CRC^28^, showed a decreasing trend in comparisons including L vs. N (FC = 0.38, p = 3.08e-02), H vs. N (FC = 0.055, p = 1.14e-04), and C vs. N (FC = 0.072, p = 1.14e-04). These findings indicated the capability of the workflow used in the study to dissect the heterogeneity of different malignant changes within the same tissue slice.

To study the variations across tissues and patients, we calculated the coefficient variations of protein abundance between three consecutive slides of the same subtype (intra-patient variation) and among all punches within the same subtype (intra-subtype variation). The results showed that, across all subtypes, the CV values for intra-patient variation were higher than those for intra-subtype variation (Figure 5A). To explore the spatial relevant molecular features, we employed a supervised random forest-based machine learning model to investigate the dysregulated proteins that may define various disease subgroups. These subgroups were further classified into four clusters (Figure 5B). UMAP visualizations of all subtype samples from all patients, based on features obtained from the random forest machine-learning model, were displayed with arrows connecting the centroid of each subtype (Figure 5C). In the Sankey plot, our results showed the significance of mitochondrial dysfunctions related to metabolism (Figure 5D), which could have crucial implications for the discovery of therapeutic targets.

**Figure 5.**
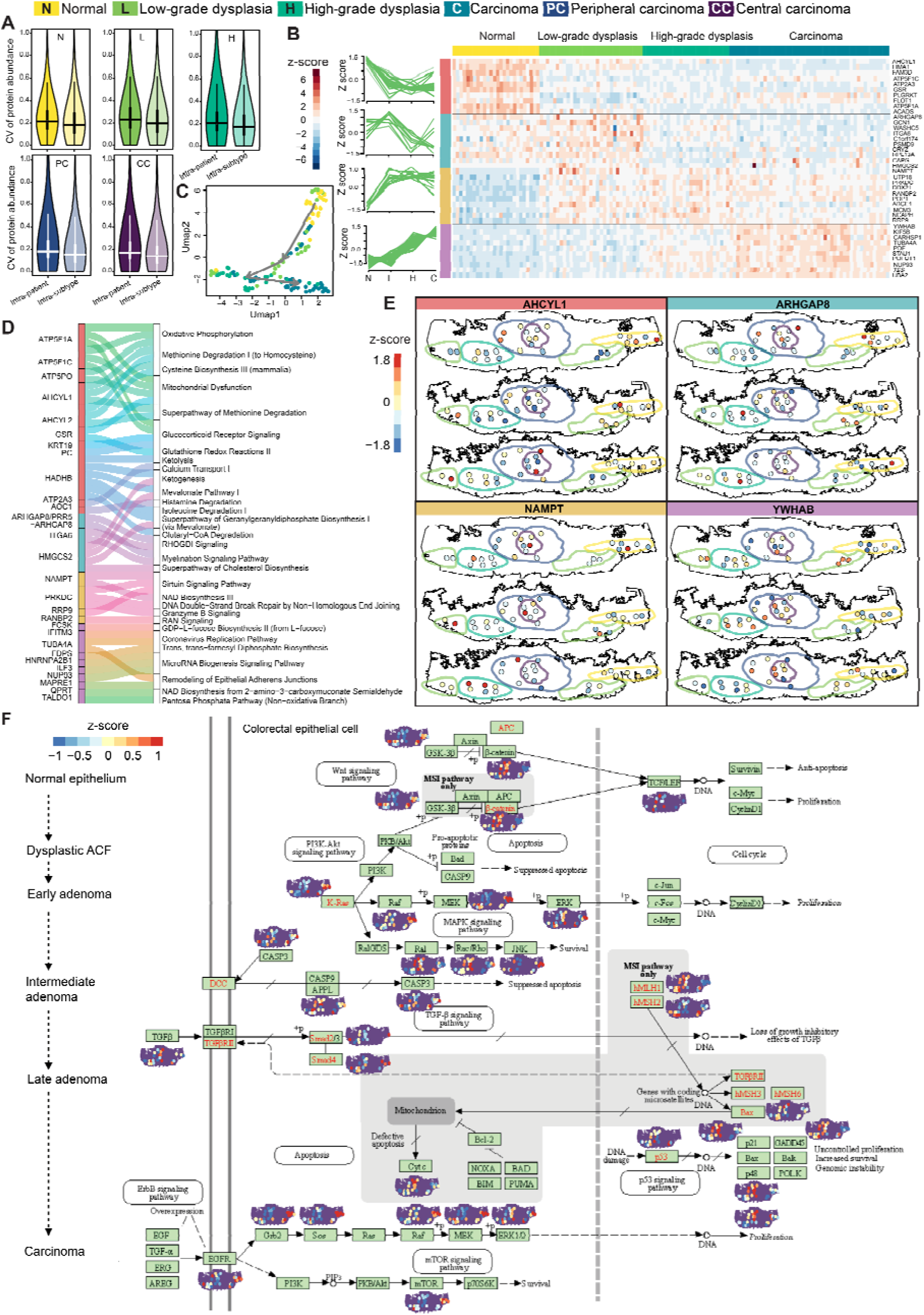
Spatial analysis of colorectal proteome expression. (A) Coefficient variations of protein abundance are depicted between three consecutive slides of the same subtype (intra-patient variation) and among all punches within the same subtype (intra-subtype variation). (B) Featured proteins, identified using a random forest-based machine learning model, are subjected to clustering using the Mfuzz algorithm. Heatmaps display the top 10 featured proteins in each cluster. (C) UMAP visualizations of all subtype samples from all patients by the features obtained from the random forest machine-learning model, with arrows connecting the centroid of each subtype. (D) A Sankey plot illustrates the correlation between enriched pathways obtained from IPA and proteins grouped based on Mfuzz clusters. (E) Spatial z-scored visualizations exhibit the expression patterns among three consecutive slides of the first patient, using the most important feature in each cluster as shown in (B). (F) Colorectal distribution maps present key proteins with average expression patterns of the three consecutive slides of the first patient involved in the KEGG pathway of colorectal cancer.

Furthermore, we examined the spatial expressions of AHCYL1, ARHGAP8, NAMPT, and YWHAB as the top feature in each cluster shown in Figure 5B, observing their heterogeneous distribution across different spatial regions and slices (Figure 5E), as well as the key cancer regulator KRAS^29^. Moreover, pathway-centric analysis unveiled differential spatial expression of multiple proteins within CRC-relevant Kyoto Encyclopedia of Genes and Genomes (KEGG) pathways (pathway entry: hsa05210) (Figure 5F). The distribution map provides insights into trends in protein expression across different stages. In the carcinoma stage, we observed a noticeable increase in the number of upregulated proteins involved in the p53 signaling pathway within the carcinoma area. Conversely, the ErbB signaling pathway exhibited a higher count of downregulated proteins in the same carcinoma area. These findings underscore the dynamic changes occurring in CRC samples during disease progression.

## Discussion

Establishing the FAXP workflow represents a significant advancement in improving lateral/volumetric resolution and enhancing the throughput for spatially resolved proteomics analysis of biological tissues. We have also optimized the hydrogel recipe to enable effective expansion proteomics analysis of FFPE samples. Key novel or optimized steps were introduced to significantly expedite the processing time and depth of the spatial proteome (Figure 6A). These include: i) slicing FFPE samples onto customized glass slides with chamber or commercial glass slides for all the scenarios of FFPE sample placement; ii) performing dewaxing and rehydration; iii) introducing autoclave-based tissue homogenization to utilize high pressure and heat to accelerate the tissue homogenization; iv) conducting reduction/alkylation on the entire tissue slice embedded in hydrogel to reduce the hands-on steps during proteomics sample preparation and increase the reproducibility; v) performing fluorescent staining or optimizing Coomassie Brilliant Blue R-250 staining after tissue expansion for quicker staining and easier de-staining; vi) facilitating manual microdissection down to 73 μm in diameter or laser capture microdissection down to subcellular organelle for higher resolution and employing proteomics sample preparation through the developed filter-aided sample preparation to reduce sample loss and increase sample processing throughput; vii) utilizing DIA-MS with timsTOF Pro and performing protein identification using a hybrid search library strategy to ensure proteome coverage. In addition, the workflow is partially automated with robotics.

**Figure 6.**
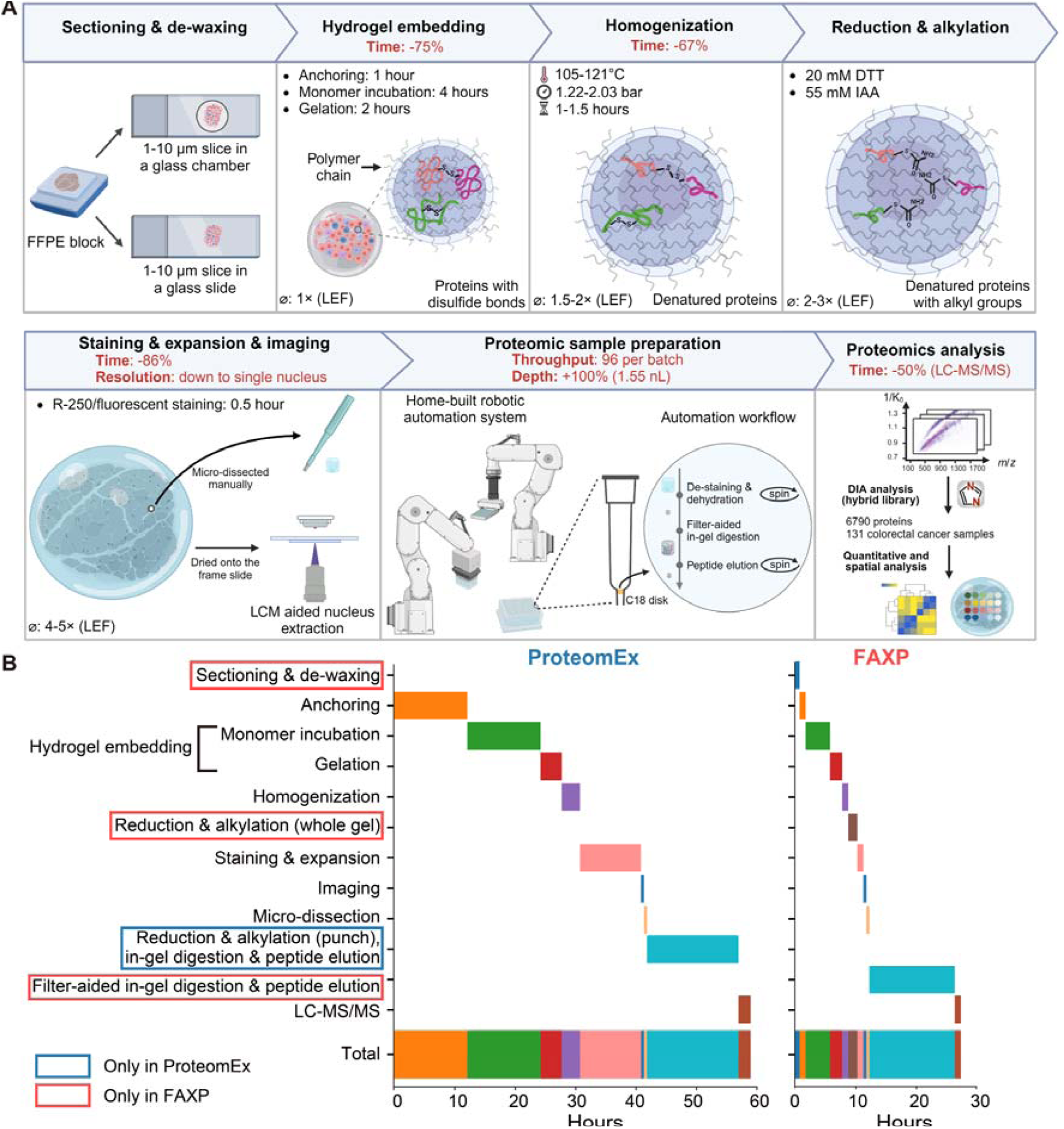
Comparative analysis of FAXP and ProteomEx. (A) Workflow of FAXP. The FAXP workflow begins with the sectioning and de-waxing of FFPE tissue block. Subsequently, the samples undergo treatment with a chemical anchor, followed by embedding into a hydrogel and homogenization using an autoclave. Either the updated Coomassie brilliant blue staining or fluorescent staining happen after reduction and alkylation on the whole-gel level, with expansion and imaging steps. After imaging, the expanded samples are micro-dissected manually or cut through laser capture microdissection (LCM) system, and peptides are collected through filter-aided in-gel digestion under the home-built robotic automation system for DIA-mass spectrometry (MS) data acquisition. For the human FFPE colorectal cancer samples, the MS-based proteomic data is searched in library-free mode using DIA-NN, generating a predicted library that is later combined with a pan-human library for a library-based search, which results in a protein matrix for downstream analysis. The spatial proteome expression data are further analyzed for spatial analysis. Notably, significant performance enhancements achieved in specific steps of this workflow are indicated in red below those steps (LEF - linear expansion factor). (B) Comparison of processing timeline. The bars represent the total duration time of each step under different workflows. Blue rectangles denote steps exclusively used under the ProteomEx, while red rectangles correspond to steps used under FAXP. Hands-on time is not displayed, as both the ProteomEx and FAXP workflows involve similar hands-on time when dealing with small sample amounts. Besides, ProteomEx involves reduction and alkylation on individual punches during conventional in-gel digestion, while FAXP performs these steps on the entire gel without further reduction and alkylation during filter-aided in-gel digestion. The faster processing time and improved workflow of FAXP are achieved from shorter duration time for various steps compared to ProteomEx. The data in FAXP are based on the processing of 5 μm FFPE mouse liver.

FAXP has achieved two remarkable features in spatial resolution and sample processing throughput. This lateral resolution is 2.2 times higher than that achieved with ProteomEx. FAXP facilitates the processing of micro-dissected samples in batches of up to 96, with the simultaneous processing of multiple batches. Collectively, FAXP has reduced the total processing time by up to 53.5% compared to ProteomEx (Figure 6B). In ProteomEx, the sample loss rate during the peptide extraction step is around 6%^14^. The FAXP method minimizes both sample and peptide loss due to fewer hands-on steps and the peptide-trapping capability of the hydrogel during sample preparation.

Another key distinction of FAXP is its adaptability to varying sample types. For instance, samples rich in extracellular matrix, such as CRC tissues^30,31^, require more rigorous homogenization conditions involving higher SDS concentration, elevated homogenization temperature, and longer processing time. This dynamic optimization, as demonstrated by different conditions in Figure S7A (Table S1), ensures that FAXP remains versatile and effective across diverse samples, highlighting its flexibility for clinical applications.

When considering CRC applications, disparities emerge among N, L, H, C, CC vs. PC subtypes in contrast to the global proteome, underlining the significance of analyzing distinct CRC subtypes to gain deeper insights into the disease’s molecular mechanisms^32,33^. Especially, FAXP exhibits the capability to discriminate heterogeneity within inter-patient, inter-slide, and intra-tissue clinical FFPE samples. This attribute bears particular importance in clinical research, enabling precise analysis and characterization of samples, thereby advancing comprehension of diseases and potential therapeutic avenues.

There are also some limitations for FAXP workflow. Optimization of the gel expansion technique for tissues rich in collagen and extracellular matrix components represents one of the challenges. Another limitation is the staining method, which currently only uses Coomassie Brilliant Blue. This falls short compared to more advanced staining methods like immunofluorescence, which is important for precise clinical observations. To enhance its usefulness in addressing heterogeneity in clinical samples, the workflow should also be able to handle samples with irregular shapes. While FAXP does improve spatial resolution compared to ProteomEx, it still cannot achieve single-cell resolution using manual operation same as LCM technique due to limitations in sample collection method, while alternatively LCM based technique can be incorporated with FAXP to further extend to the single-cell and subcellular organelle resolution. In this paper, we have demonstrated the application of FAXP in analyzing the proteome of a single nuclear. More types of subcellular organelles could be potentially analyzed in future experiments.

In summary, the introduction of the FAXP workflow represents a significant step forward in hydrogel-based spatial proteomics, addressing challenges in tissue expansion, sample preparation and data analysis. Its notable strengths include increased throughput of sample preparation, better sample compatibility, improved spatial/volumetric resolution, and enhanced reproducibility.

## Materials and Methods

### Preparation of mouse organ FFPE tissue samples

All animal maintenance and experimental procedures were conducted according to the Westlake University Animal care guidelines, and all animal studies were approved by the Institutional Animal Care and Use Committee (IACUC) of Westlake University, Hangzhou, China, under animal protocol #22-083-GTN.

Male C57BL/6J mice (four-month old) were anesthetized deeply using 1% sodium pentobarbital to perform the procedures. Subsequently, transcardial perfusion was performed using 1× phosphate-buffered saline (PBS) and 4% paraformaldehyde (PFA; Electron Microscopy Sciences, the United States). The mouse liver and kidney were dissected and postfixed in 4% PFA for 6 hours at 4°C. Fixed tissues were dehydrated using increasing ethanol concentrations: 75%, 95%, and 100% for 30 minutes each. Next, the dehydrated tissues underwent infiltration with paraffin wax at 60°C. The embedded tissues were sliced into either 5-µm or 10-µm in thickness, utilizing a rotary microtome (Leica RM2255, Germany) according to a standard protocol and mounted on the glass slides.

### Collection of FFPE samples from patients

Archival FFPE colorectal tissue samples were acquired from three patients diagnosed with CRC at the Second Affiliated Hospital of Zhejiang University School of Medicine, Hangzhou, China. The samples were sliced to a thickness of 8 μm with a rotary microtome (Leica RM2255, Germany). The tumor grading followed the Tumor, Node, Metastasis (TNM) classification system of colorectal cancer, as illustrated in Figure 4. Informed consent was obtained from all patients prior to their surgeries. Patient-derived tissues were embedded in paraffin according to the approved institutional review board protocol at the same hospital. Human tissue samples were collected with the approval of the Institutional Ethics Committee of the Second Affiliated Hospital of Zhejiang University School of Medicine (No. 2020-322). The study was also approved by the ethics committee of Westlake University (Permission number: 20220913GTN001).

### Tissue expansion and staining

The chemical composition and storage conditions for all reagents and buffers used for tissue expansion, staining, and sample preparation are described in Table S4. Suppliers and catalog numbers for chemicals, reagents, and accessories used for FAXP are listed in Table S5. The paraffin-embedded sections were mounted onto glass slides and subjected to deparaffinization. Two rounds of 10-minute heptane treatments were performed. The sections then underwent a rehydration procedure involving decreasing ethanol concentrations: 100%, 90%, and 75% for 5 minutes each followed by rehydration in ddH_2_O for 5 minutes. The rehydrated FFPE tissue sections were then ready for tissue expansion. The rehydrated CRC tissue sections were subjected to H&E staining following the manufacturer’s protocol (Shanghai Yuanye Bio-Technology, China).

The tissue expansion procedure followed the previously described protocol^14^, incorporating certain modification. Briefly, rehydrated tissue sections were rinsed in 1× PBS and treated with protein anchoring solution for either 1 hour or 2 hours at 25°C. Protein anchoring time was reduced compared to the original protocol because the thinner tissue sections were used in this study (5-µm or 10-µm vs. 30-µm). The anchoring solution was removed by washing the samples three times with the anchoring termination buffer for 5 minutes each. The NSA-anchored tissue was incubated with activated monomer solution in a gelation chamber at 4°C to infuse monomers. Subsequently, the treated tissue was transferred to a vacuum oven for polymerization reaction at 37°C in a nitrogen gas atmosphere for 2-3 hours. To accelerate homogenization step, we decided to use sample autoclavation similarly to dExPath protocol^34^. The formed tissue-hydrogel composite was transferred to a dish with the protein denaturation buffer. The buffer was adapted from the dExPath protocol^34^, comprised of 50 mM tris(hydroxymethyl)aminomethane (Tris), 20% sodium dodecyl sulfate (SDS), 25 mM ethylenediaminetetraacetic acid, disodium salt, dihydrate (EDTA-Na_2_·2H_2_O), and 200 mM NaCl. The homogenization condition involved a temperature range of 105-121°C maintained under 1.22-2.03 bars for 60-90 minutes in the autoclave. To improve mechanical stability of the hydrogel after autoclaving, we increased cross-linker concentration 17.5-fold from original (Table S1). Homogenized samples were transferred to 9 cm petri dishes and washed with 1× PBS three times. Reduction and alkylation of proteins were performed by adding the reduction buffer (20 mM dithiothreitol (DTT) in 1× PBS) and alkylation buffer (55 mM iodoacetamide (IAA) in 1× PBS) and incubating for 30 minutes, respectively. The samples were washed with 1× PBS three times for 10LJminutes each time. The hydrogel sample was incubated in ddH_2_O and allowed to expand to its maximum size. We found out that using Coomassie Brilliant Blue R-250 (Sangon, China) staining allowed us to stain and expand samples in less than 1 hour compared to the almost 10 hours of staining and expansion in the original protocol. Imaging was performed using Zeiss Fluorescence Stereo Zoom Microscope (Axio Zoom.V16) in brightfield mode, controlled by ZEN 3.1 software. Furthermore, we found DAPI staining was also compatible with our workflow (Figure S8). By using hydrogel samples with a diameter of 488 µm derived from mouse liver slices, we could identify an average of 830 peptides and 350 proteins using 0.5-hour DIA-MS by timsTOF Pro.

### Optimization of reduction and alkylation conditions

Three hydrogels of mouse liver FFPE slices (10 μm) were employed to determine the best conditions for protein reduction and alkylation on whole tissue slice samples. Four sets of reduction and alkylation conditions (10 mM DTT and 55 mM IAA; 20 mM DTT and 55 mM IAA; 20 mM TCEP and 55 mM IAA; 45 mM TCEP and 100 mM IAA) were then tested separately on the gels (as shown in Figure 1H). The combination of 20 mM DTT/55 mM IAA resulted in the highest carbamidomethyl modification rate close to an average 98% (Figure 1I). This condition exhibited comparable rates of cysteine-containing peptide and protein identifications when compared to the ProteomEx method (indicated No. 1 in Figure 1I), while also demonstrating superior peptide identification performance.

### Proteomic sample preparation using conventional in-gel digestion

Manual microdissections were performed on the expanded and Coomassie-stained tissue-hydrogels (following reduction and alkylation). The excised tissue-hydrogel samples were de-stained in 50% acetonitrile (ACN) and 50% ddH_2_O for 30 minutes at 30°C. Samples were then dehydrated twice in a solution containing 50% ACN and 100 mM ammonium bicarbonate (ABB) for 10 minutes each and dried in a SpeedVac at 45°C for 5 minutes. Protein digestion was done using 12.5 ng/µL trypsin (Hualishi Tech. Ltd, China) in 10% ACN and 90% 50 mM ABB. The digestion process occurred at 37°C for 4 hours, followed by overnight incubation with an extra 10 mM ABB.

The supernatant of the digested peptide solutions was collected first. After that, 100 µL of 25 mM ABB was added to collect the resulting supernatant. Then, 100 µL of 50% ACN and 2.5% formic acid (FA) was added and vortexed for 10 minutes; this step was repeated three times with the supernatant collection each time. Next, 100 µL of 100% ACN was added until the hydrogel fragments became small and white. The peptide solutions were then concentrated using a SpeedVac to 20-30 µL. Purification was done using C18 micro spin columns (Thermo Fisher Scientific, the United States). After purification, the samples were dried with a SpeedVac, stored at -80°C, and ready for later LC-MS/MS analysis.

### Proteomic sample preparation using filter-aided in-gel digestion

One plug of the C18 disk (Empore, the United States) was assembled into either the 10 µL or the 200 µL tip to form a spintip device. The spintip device was first rinsed and activated with 20 µL or 100 µL of 80% ACN for tightening the C18 pad. 10 or 50 µL of 50% ACN and 50% 100 mM ABB were added. After transferring the excised tissue-hydrogel samples, with sizes based on experimental needs, into the devices, another 10 or 50 µL of 50% ACN and 50% 100 mM ABB were added and incubated for 15 minutes. The samples were dehydrated by adding 20 µL or 100 µL of 100% ACN until they became small and white and then centrifuged at 150 g for 2.5 minutes. Protein digestion was performed using 12.5 ng/µL trypsin (Hualishi Tech. Ltd, China) in 10% ACN and 90% 50 mM ABB. The digestion process took place at 37°C for 4 hours, followed by overnight incubation with an extra 10 mM ABB.

The digested peptide solutions were collected and combined through the following steps: Firstly, 6 µL or 12 µL of 2% ACN and 0.1% trifluoroacetic acid (TFA) were added and incubated for 15 minutes, followed by centrifugation at 150 g for 3 minutes. Secondly, elution was performed with 6 µL or 20 µL of 70% ACN and 0.1% TFA, with two 15-minute incubations and subsequent centrifugation at 150 g for 5 minutes after each incubation. Thirdly, elution was performed using 6 µL or 20 µL of 100% ACN, followed by incubation for 15 minutes and then centrifugation at 150 g for 5 minutes. The resulting peptide solutions were concentrated to dryness using a SpeedVac and stored at -80°C, ready for later LC-MS/MS analysis.

### Determination of detection range of the minimal volume

Tissue volumes were estimated using commercially available biopsy punches based on needs (0.35 mm, 0.5 mm, 0.75 mm, 1 mm, 1.5 mm, 2 mm, 3 mm; Integra Miltex, the United States). Specifically, the formula to calculate the volume is: volume (nL) = π × (D/2F)^2^ × h × 10^-6^, where D is the diameter (µm) of the punch, F is the expansion factor calculated as the square root of the ratio of the tissue area after expansion to the area before expansion, and h is the thickness (µm) of the slice. The detection range calibration curves were estimated using Local Polynomial Regression Fitting estimation (loess) using the number of identified peptides and proteins.

### Single nucleus collection, sample preparation and proteomics analysis

A hydrogel sample of 5 μm FFPE mouse liver was stained through SYPRO Ruby protein gel stain (Thermo Fisher Scientific, the United States) for 30 minutes. Then the hydrogel sample was attached onto a polyester (POL) frame slide (Leica, Germany). Single nuclei were individually imaged and cut by a MMI CellCut (Molecular Machines & Industries, Germany) and collected by diffuser caps (Molecular Machines & Industries, Germany). Collected samples were then processed through 10 µL tip under filter-aided in-gel digestion. Peptides were analyzed through an Orbitrap Astral mass spectrometer (Thermo Fisher Scientific, the United States) using DIA mode under the liquid chromatography conditions and MS parameters suggested by the Thermo Fisher Scientific (technical note: 002255).

### Liquid chromatography

The peptide samples were separated on a 15 cm × 75 μm silica column custom packed with 1.9 μm 100 Å C18 aqua installed into nanoElute® system (Bruker Daltonics, Germany). The mobile phase was mixed with buffer A (MS grade H_2_O + 0.1% FA) and buffer B (MS grade ACN + 0.1% FA). The flow rate was set at 300 nL/min. For both 30-minute and 60-minute linear LC gradients, the buffer B (%) was linearly increased from 5% to 27% in 25 minutes and 50 minutes separately, followed by an increase to 40% within 5 minutes and 10 minutes separately, and a further boost to 80%. The results showed that 1-hour LC gradient resulted in more peptides and proteins, along with an increase in the mean peak full width at half maximum (FWHM) from an approximate average of 2.5 to 3.5 (Figure S7B).

### Proteomic data acquisition using MS

Eluted peptides were analyzed in a trapped ion mobility spectrometry (TIMS) combined with a quadrupole time-of-flight mass spectrometer (timsTOF Pro, Bruker Daltonics, Germany) via a CaptiveSpray nano-electrospray ion source. MS data were acquired using the Bruker Otofcontrol v6.2 and HyStar v5.1 mass spectrometer control software.

Data-dependent acquisition (DDA) was performed in the PASEF mode with 10 PASEF scans per topN acquisition cycle. The ramp time was set at 100 ms, achieving a total cycle time of 1.17 s. The ion mobility was scanned from 0.7 to 1.3 Vs/cm^2^. MS1 and MS2 acquisition was performed in the m/z range from 100 to 1700 Th. Precursors that reached a target intensity of 20,000 cts/s were dynamically excluded for 0.4 minute. Singly charged precursors were excluded by their position in the m/z-ion mobility plane. For the data-independent acquisition Parallel Accumulation Serial Fragmentation (diaPASEF) mode, the majority of settings were the same as in DDA mode, except the ramp time was set at 166 ms. We defined the isolation windows as shown in Table S3. For the determination of the detection range, both DDA and DIA methods were used for sample collection. A total of 33 DDA files and 33 DIA files were collected for evaluation purposes.

### Protein identification and quantification

DDA data were analyzed using FragPipe (version 18.0) platform^35,36^ with the MSFragger (version 3.5) search engine against a FASTA file from SwissProt containing 16,985 mouse protein entries (downloaded in September 2018) and 16,985 decoy sequences for mouse samples and a FASTA file from SwissProt containing 20,386 human protein entries (downloaded in November 2022) and 20,386 decoy sequences. The “IM-MS” was selected as the MS data type under “Input LC-MS Files” section. A fragment mass tolerance was set at 0.05 Da. The enzyme used for digestion was trypsin, with cleavage occurring after “KR” residues but not preceding “P” residues. To evaluate the cysteine modification rates, carbamidomethylation was considered variable modification, while in the other DDA files, it was set as fixed.

DIA data analysis was performed by DIA-NN (version 1.8.1)^37,38^. The false discovery rate for precursors was set at 1%. Fixed modifications included “C carbamidomethylation,” while variable modifications included “N-term M excision” and “Methionine oxidation.” The peptide length range was set between 7 and 50. The precursor charge range was set between 2 and 4. Both m/z ranges for precursor and fragment ions were set between 100 and 1700. “Unrelated runs,” “Use isotopologues,” and “MBR” options were selected. Heuristic protein inference was chosen under mouse liver-based studies. Protein inference was set as “off,” and the quantification strategy was set as “Robust LC (high precision).” Other settings remained as default values. The DIA data searching employed an in-house generated library for mouse liver-based studies. For the single-nucleus studies, together with the data from 0.042 nL of In-tip DIA-MS, proteomic data was searched using library-free mode by DIA-NN. Similar as the settings shown above, the modifications were mainly made for precursor (300-1800) and fragment ion (200-1800) m/z ranges. For the CRC application, proteomic data was searched using library-free mode by DIA-NN. This generated a predicted library, which was then merged with an in-house generated pan-human library to form a hybrid library. Only the precursors do not present in the pan-human library were used in combining the spectral library. The hybrid library was then used for a library-based search. The peptide and protein numbers in the DDA analysis were extracted from the FragPipe peptide.tsv and protein.tsv files, whereas in the DIA analysis, they were acquired from the stripped sequences in DIA-NN pr.matrix and unique proteins in the pg.matrix.

### Batch design and data analysis for MS acquisition

The MS batches were systematically arranged, balanced, and randomized, considering patient ID, slide ID, region, and replicate number features (Figure S6A). This resulted in acquiring six batches, with 22 injections per batch. It’s worth noting that one file failed due to an MS bug during acquisition, resulting in a total of 131 files. The coefficient of variation and Pearson correlation analysis for global precursors and proteins demonstrated robust reproducibility in the MS data (Figure S6B). When examining pooled samples across different batches, we observed high consistency with pool samples (Figure S6C), and the missed cleavage rates exhibited low and consistent values (Figure S6D). We evaluated five distinct spectral searching conditions aimed at achieving deepest identification performance, including a colon library developed in-house with default DIA-NN parameters and optimal DIA-NN parameters for mass range (which was also optimized for retention time window); a hybrid-library search approach using the combined library from predicted library obtained via a DIA-NN library-free search for those proteins not included in the spectral library; a pan-library consisting of multiple tissue types with a hybrid search approach.

### Feature selection using the random forest model

The unsupervised clustering approach involved selecting the protein expression matrix derived from each patient sample. The k-means method was then applied for clustering spatial locations using the proteome abundance into four clusters for each slice, utilizing different shapes to represent the cluster category on a location map and colors to indicate the subtype. Finally, the central point of each cluster was calculated to generate a disease process pathway map. We used a random forest-based model to select the top 100 features that separated four disease subgroups (normal, low-grade dysplasia, high-grade dysplasia, carcinoma) using 70% of samples as the training set and 30% of total samples as the test set. This approach yielded an accuracy of 81% and an AUC of 0.967. These selected features were then analyzed with Fuzzy c-means soft clustering (provided by mfuzz packages) into four clusters. Proteins upregulated in each disease subgroup were visualized as network modules, with proportions of each cluster indicated as pie charts.

### Spatial visualization of protein expression

With the mapped locations of punches based on microscopic images, each protein was visualized as a z-score using scaled colors, where subgroup boundaries were determined manually. The pathway visualization highlighted the proteins within the KEGG colorectal cancer pathway (pathway entry: hsa05210).

### Statistical analysis

Unpaired Welch’s t-test was performed for all number comparisons, and statistical visualizations were drawn in PRISM (10.0.3) and R (version 4.2.2). Isotropic analysis calculations were performed using MATLAB (version R2021a), following the previously published protocol^14^. The differentially expressed protein comparison was performed at a threshold of adjusted P-value <0.05 and fold change >2 (P-values were adjusted by the method of Benjamini–Hochberg). GO term and pathway enrichment analyses were performed by ingenuity pathway analysis (IPA), in which a right-tailed Fisher’s exact test estimated the P-value.

As to the single nucleus analysis, the protein matrix from DIA-NN was transformed using a logarithmic scale and then normalized across samples using quantile normalization. The Pearson correlation coefficient was calculated after the removal of missing values. Unless otherwise specified, missing values were imputed by using the minimum value. Between-group differences were evaluated using Welch’s t-test, and the P values were adjusted using the Benjamini–Hochberg procedure in R (version 4.4.0). Gene ID conversion and Gene Ontology (GO) enrichment analysis were executed using the clusterProfiler (version 4.11.0), with all default options except for the maxGSSize which was set to 2500. The annotation of GO terms was performed using org.Mm.eg.db (version 3.18.0). Redundant GO terms were simplified using the default options of the clusterProfiler::simplify function. The Reactome pathway-based analysis was conducted using the ReactomePA (version 1.47.0) package.

### Data availability

The mass spectrometry proteomics data have been deposited to the ProteomeXchange Consortium via the PRIDE^39^ partner repository with the dataset identifier PXD041412.

## Acknowledgments

We thank the NSFC grant (Grant No. 92259201 to Dr Tiannan Guo), the National Key R&D Program of China (Grant No. 2022YF0608403 to Dr Yi Zhu) and the Key Research and Development Program of Zhejiang Province (Grant No. 2022C03037 to Dr Tiannan Guo). Figure 6A was created with Biorender.com. We thank Cuiji Sun for teaching gel-making approaches based on ProteomEx, and Liang Yue for sharing home-built DIA libraries. We thank the Research Center for Industries of the Future (RCIF) at Westlake University for partially supporting this work. Declaration of generative AI and AI-assisted technologies in the writing process during the preparation of this work the authors used ChatGPT to improve language and readability. After using this tool/service, the authors reviewed and edited the content as needed and take full responsibility for the content of the publication.

## Author contributions

T.G., Y.Z., and K.D.P conceived the project. Z.D., Y.Z., K.D.P., and C.W. developed and benchmarked the method. T.C. provided the clinical samples. Z.D., C.W., and J.C. performed the proteomic experiments. Q.X., and Z.D. participated in the robotic automation system establishment. F.Z., Z.D., and W.J. conducted proteomic data analysis and Z.D. performed the isotropic analysis. D.Z., F.Z., K.D.P. and T.G. wrote the manuscript, and the others revised the manuscript. T.G., Y.Z., and K.D.P. supervised the project.

## SUPPLEMENTAL INFORMATION

**Figure S1.**
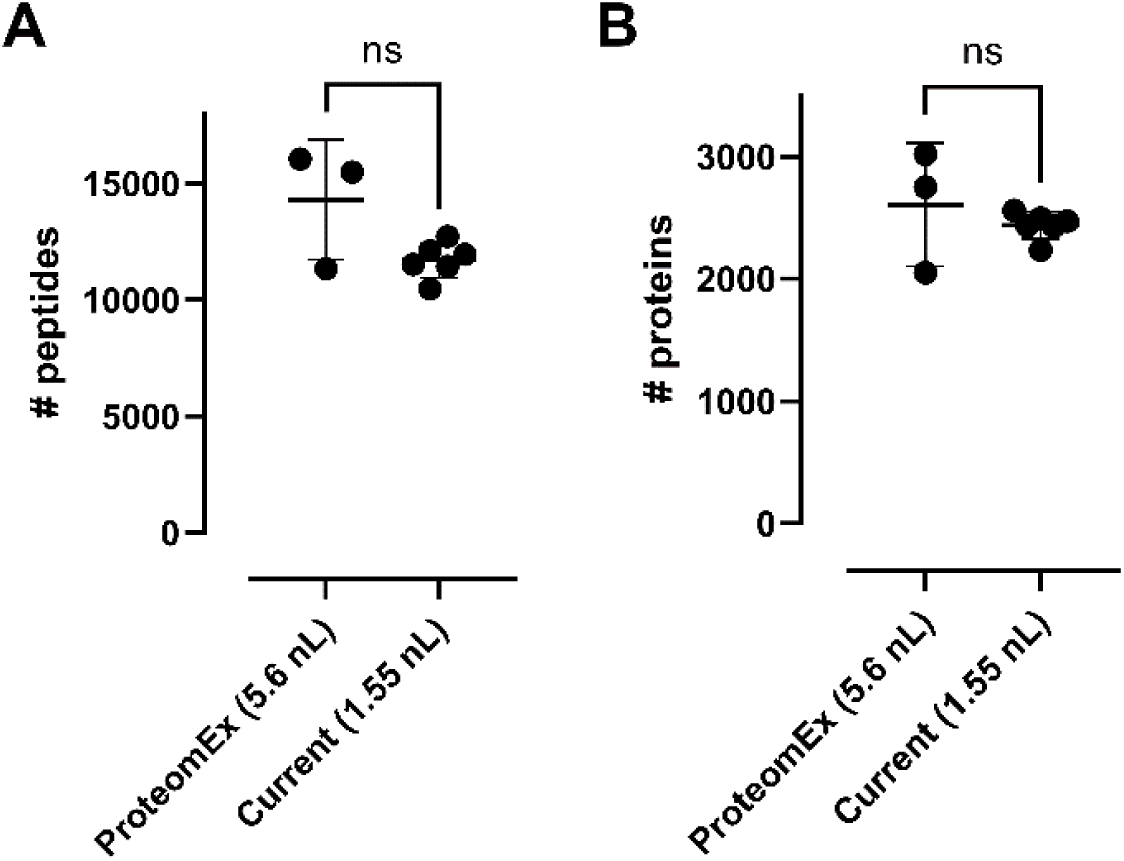
Acceleration of the ProteomEx timeline does not sacrifice proteome coverage for mouse liver tissue. (A) Peptide and (B) protein identifications of the mouse liver samples between ProteomEx (5.6 nL; LC gradient time 80 min) and the current FAXP workflow (1.55 nL; LC gradient time 60 min) using timsTOF Pro DDA mode (data for ProteomEx is from the original study^14^, n=3; data for FAXP is n=6 technical replicates from one liver slice each). All data points are presented as mean values ± SD.

**Figure S2.**
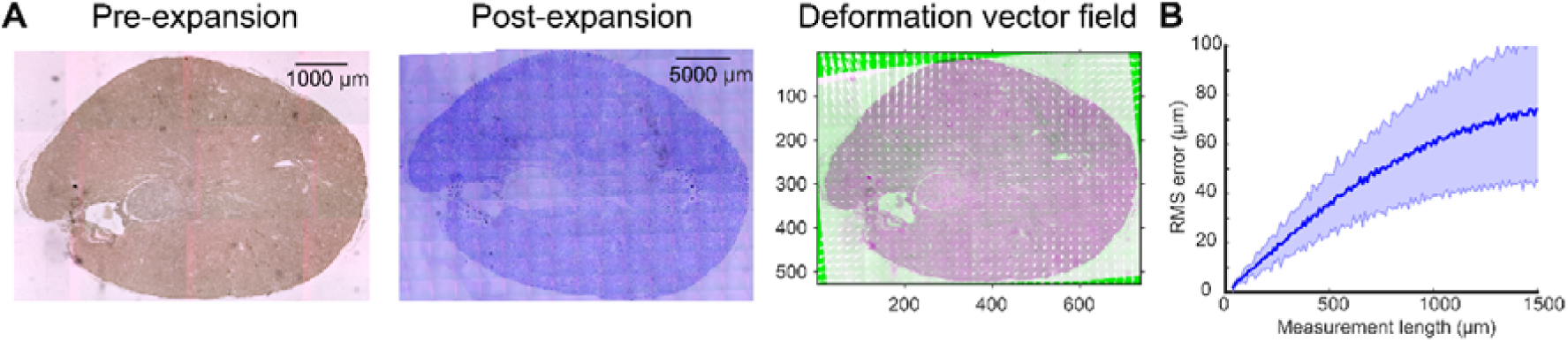
Isotropic analysis of mouse kidney slice sample expansion using FAXP under autoclave (121°C, 1.5 h). (A) Bright-field images of mouse kidney tissue slice pre-expansion (left panel) and post-expansion (middle panel; Coomassie-stained) and overlay (right panel) of pre-expansion image (magenta pseudo-color) and registered post-expansion image (green pseudo-color). White arrows represent the deformation vector field. (B) Comparison of root-mean-square (RMS) measurement length error between pre-expansion and post-expansion kidney slice images (n = 3); one example is shown in (A).

**Figure S3.**
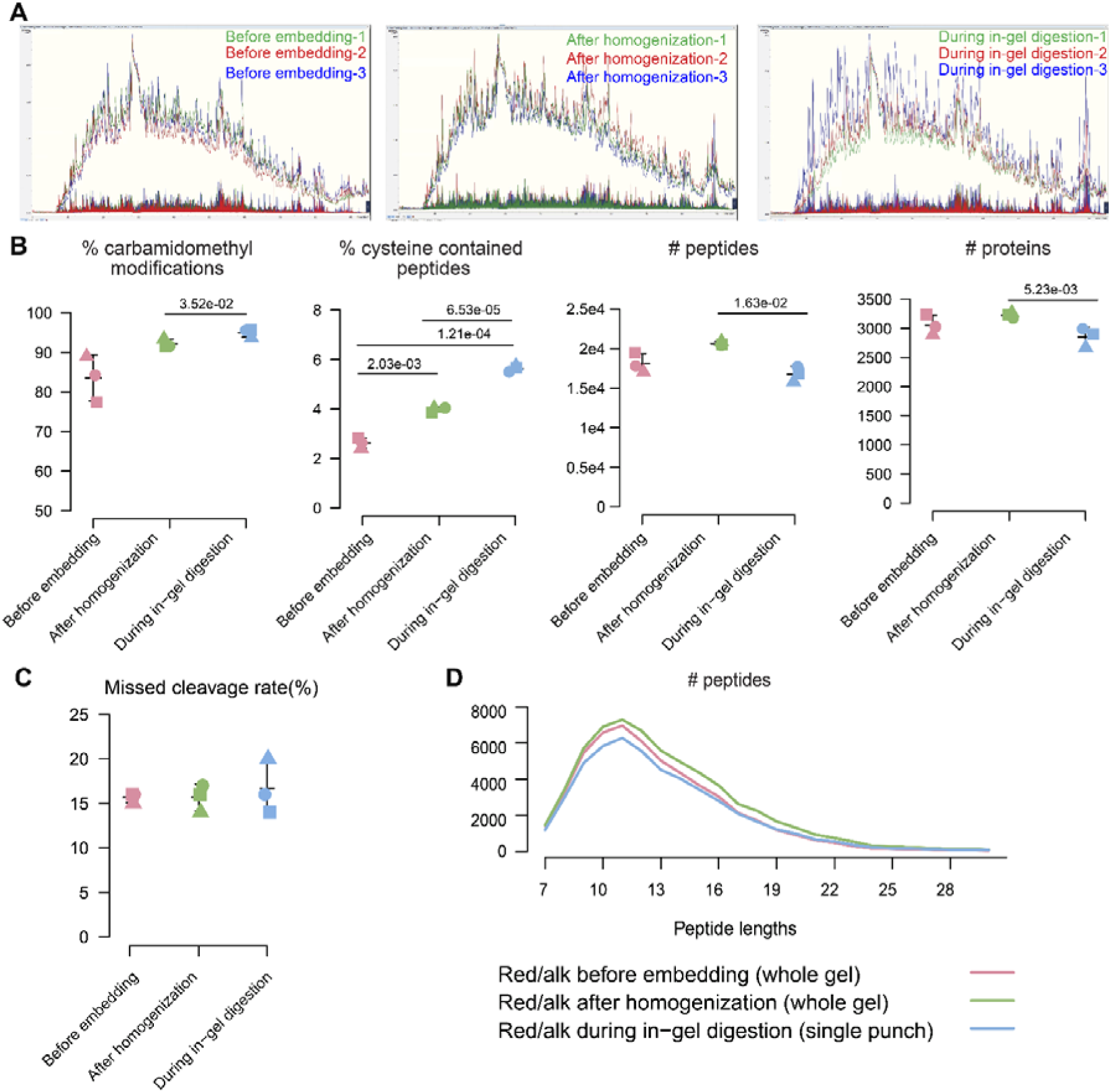
Performance of reduction and alkylation at different stages using mouse brain slices. (A) MS1 and MS2 spectra of the peptides for the samples of reduction and alkylation happens at different stages including before hydrogel embedding (whole gel), after homogenization before staining (whole gel), and during in-gel digestion (single punch). Comparison of MS1 and MS2 chromatograms across the LC retention time for the same amounts of peptides (3-mm punches) onto timsTOF Pro (90 min gradient). (B) Comparative proteomic results of the three reduction and alkylation stages, including percentages of carbamidomethyl modifications and cysteine-containing peptides, as well as the number of identified peptides and proteins. All data points are presented as mean values ± SD. (C) Missed cleavage rates of the identified peptides between three stages. (D) A curve plot shows the distribution of peptide lengths between three stages. Related to Table S2.

**Figure S4.**
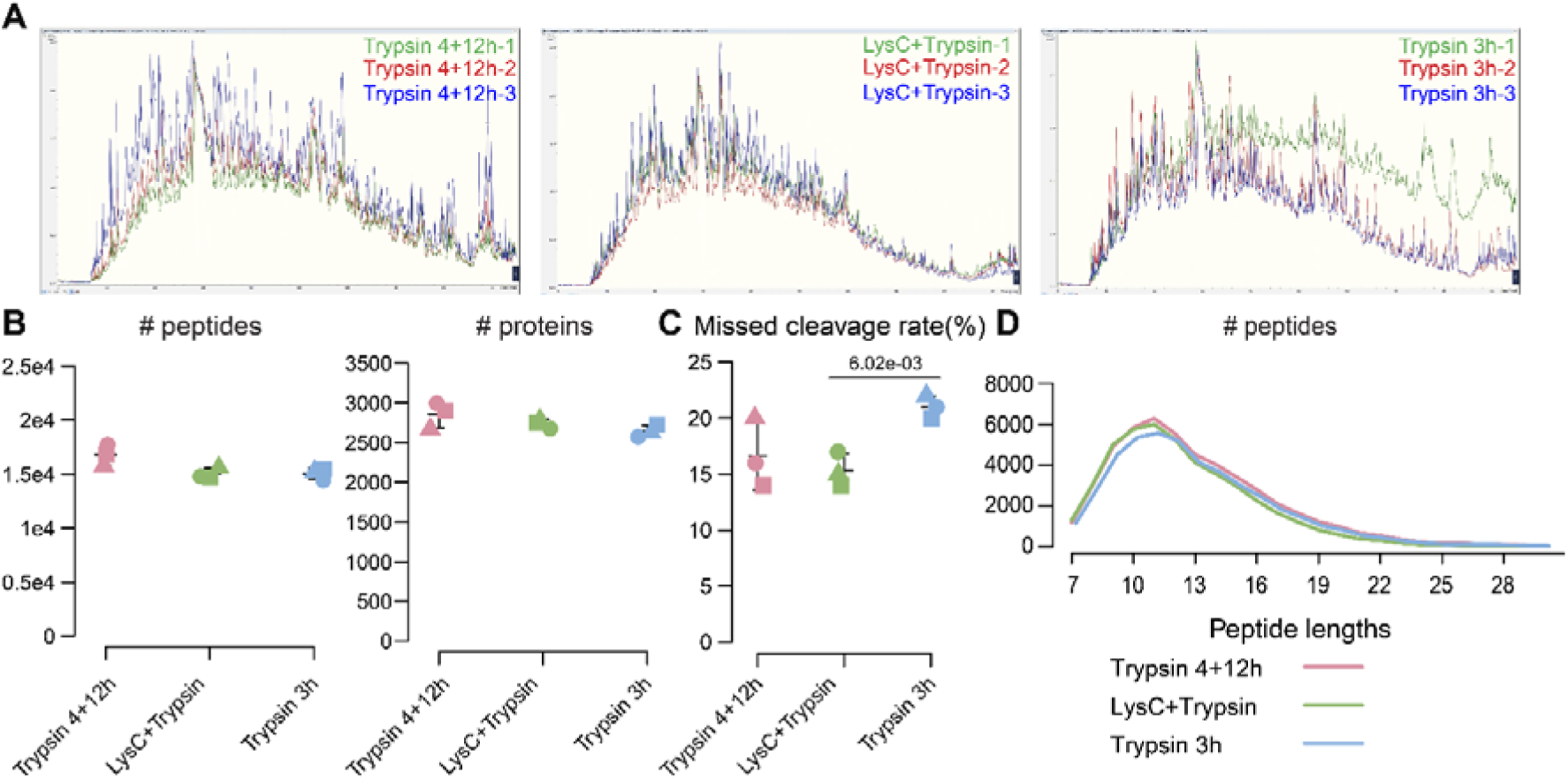
Performance of different enzyme digestion conditions using mouse brain slices. (A) MS1 and MS2 spectra of the peptides for the samples with different enzyme digestion conditions. Comparison of MS1 and MS2 chromatograms across the LC retention time for the same amounts of peptides (3-mm punches) onto timsTOF Pro (90 min gradient). (B) The number of identified peptides and proteins of the three enzyme digestion conditions. All data points are presented as mean values ± SD. (C) Missed cleavage rates of the identified peptides between three conditions. (D) A curve plot shows the distribution of peptide lengths between three conditions. Related to Table S2.

**Figure S5.**
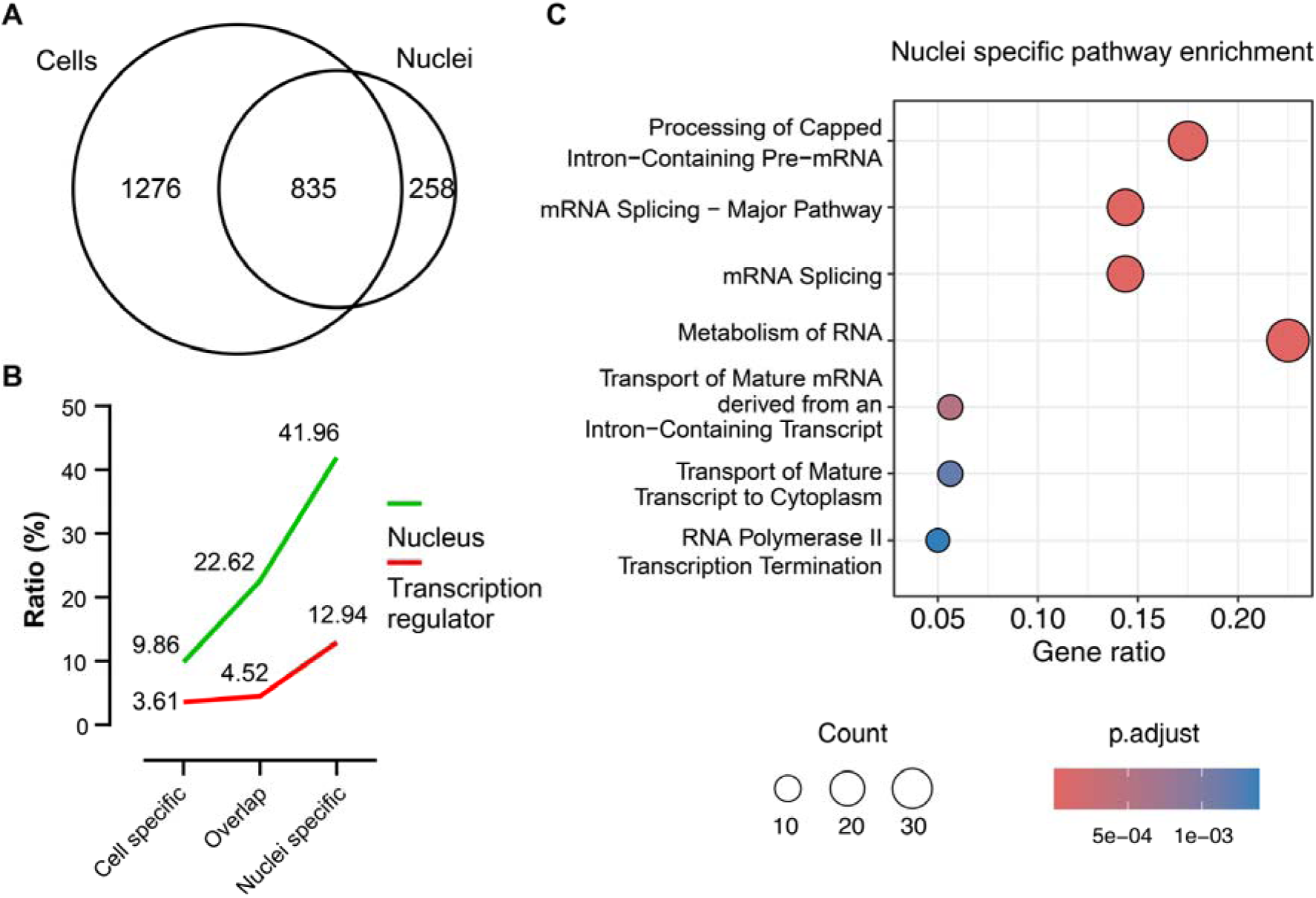
Comparative analysis between mouse hepatocytes and single nuclei. (A) Venn diagram of identified proteins for the mouse hepatocytes and single nuclei for the samples shown in Figure 3D. (B) Nucleus and transcription regulator ratio trends of the identified cell specific (1276), overlap (835) and nuclei specific (258) proteins shown in A enriched by IPA. (C) Pathway enrichment of the nuclei specific proteins.

**Figure S6.**
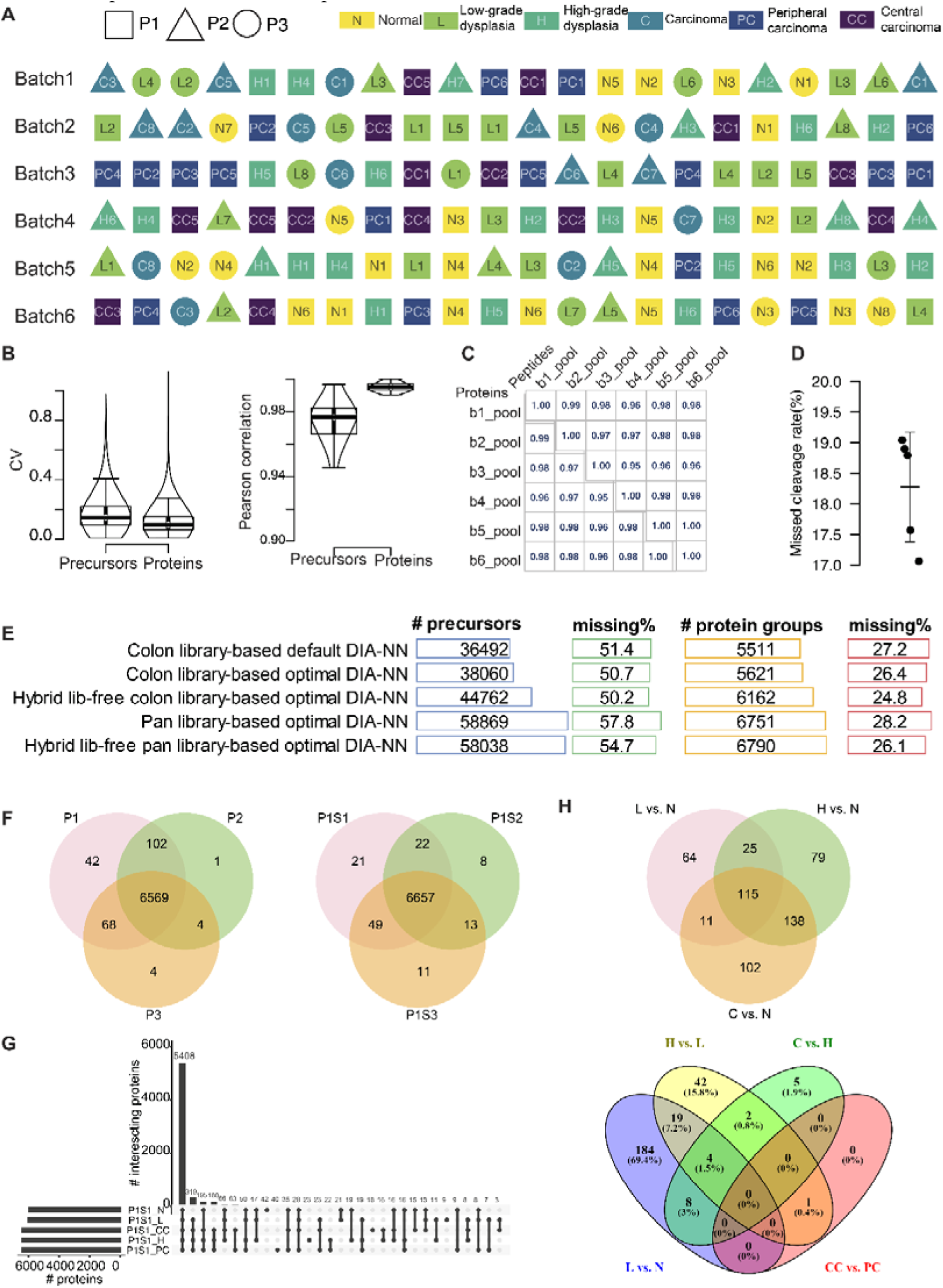
Quality control of identification results and batch design of colorectal cancer samples. (A) Batch design for the colorectal cancer samples. (B) Coefficient of variation and Pearson correlation distribution of the global precursors and proteins. (C) Pearson correlations of proteins and peptides quantification for all pooled samples from each batch. (D) Missed cleavage rates of all the pooled samples from each batch. (E) Optimizations of database search strategies. (F) Venn diagram of identified proteins for the three patients (left) and the three consecutive slides of the first patient (right). (G) Upset plots for one slide from one patient for different subtypes. (H) Venn diagrams showing the differentially expressed protein overlaps for selected comparisons.

**Figure S7.**
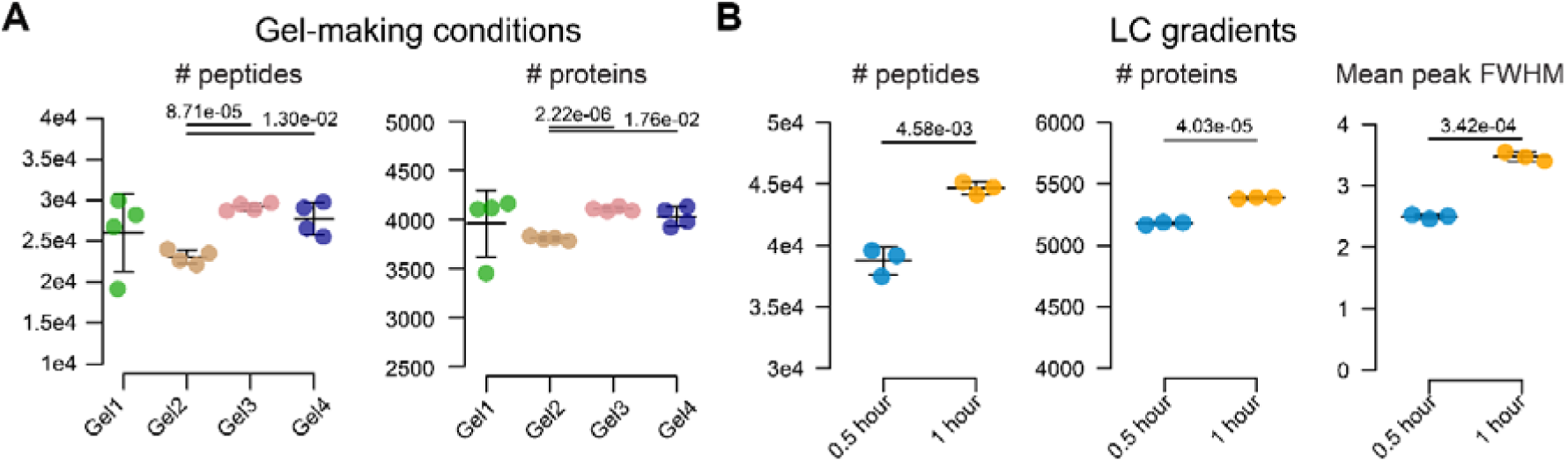
Optimization of gel-making conditions and liquid chromatography gradients. (A) Optimization of four gel preparation conditions (Table S1), involving gel compositions, homogenization buffers and autoclave conditions (n = 4). All data points are presented as mean values ± SD. (B) Liquid chromatography optimization with linear gradients of 0.5 hour and 1 hour (n = 3), reported with the number of identified peptides and proteins, as well as mean peak full width at half maximum (FWHM). All data points are presented as mean values ± SD.

**Figure S8.**
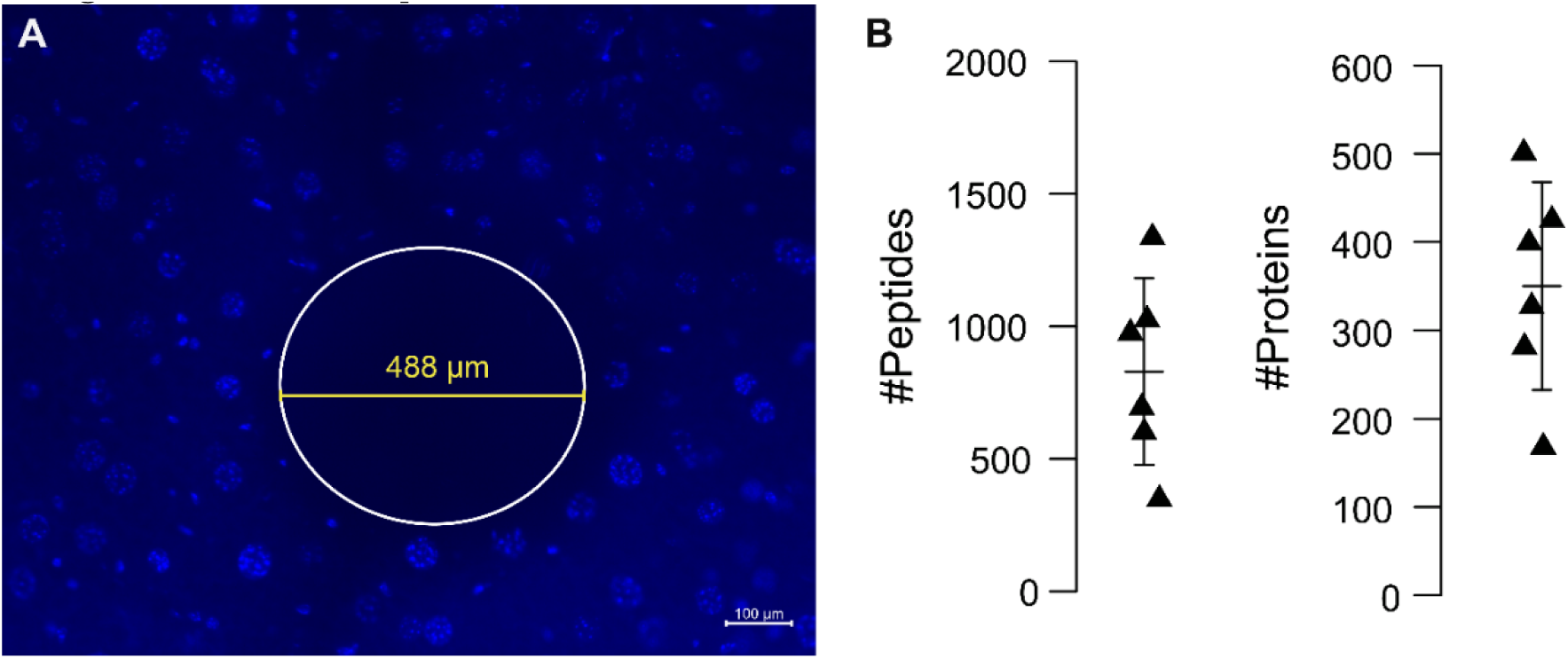
Small amount sample collection of mouse liver slices guided by DAPI staining. (A) Representative image of mouse liver slice (5 μm in thickness) stained with DAPI. After collecting punches around 488 µm in diameter, the samples were processed through filter-aided in tip sample preparation. (B) Number of peptide and protein identifications from the collected tissues guided by DAPI staining acquired by DIA-MS mode using timsTOF Pro (25 min gradient). Data are presented as mean valuesLJ±LJSD.

**Note S1.** Exploration of peptide recovery from microdissected tissue-hydrogel samples.

**Table S1.** Optimizations of gel-making conditions.

**Table S2.** Optimizations of reduction and alkylation steps as well as in-gel proteolytic digestion.

**Table S3.** MS settings for DIA-PASEF.

**Table S4.** Chemical composition of reagents and buffers for FAXP.

**Table S5.** Main chemicals, reagents, consumables, and instruments used for FAXP and their corresponding suppliers and catalog numbers.

**Supplementary Video 1.** LCM single-nucleus isolation.

