## Supplementary note 1 and supplementary table 1-5 for "Filter-aided expansion proteomics"

**Note S1. Exploration of peptide recovery from microdissected tissue-hydrogel samples.**

We started with testing whether the reduction/alkylation steps, standard treatment to increase the yield of cysteine containing peptides^1^, could be performed on the whole tissue slice either before hydrogel embedding or after tissue-hydrogel composite homogenization. Compared to the original in-gel digestion protocol, reduction and alkylation performed after homogenization step on whole tissue section resulted in slightly increased peptide and protein identifications (by 22.8% and 13.0%, respectively) and improved distribution of peptide lengths while yielding similar carbamidomethyl modification rates, cysteine-containing peptide rates, and missed cleavage rates (Figure S3). These results demonstrated that reduction and alkylation can be done on the whole tissue section samples after homogenization, reducing number of manual treatment steps during in-gel digestion streamlining the protocol.

Next, we optimized in-gel proteolytic digestion conditions utilizing 1) one round of trypsinization (3 h); 2) two rounds of trypsinization (4 h+12 h); 3) combination of LysC and trypsinization (4 h+12 h). All conditions yielded a comparable number of identified peptides and proteins (Figure S4; Table S2). However, the single-round trypsin digestion exhibited slightly higher missed cleavage rates compared to "LysC + Trypsin" digestion (Figure S4C). In conclusion, in-gel proteolytic digestion can be carried out in a wide range of conditions while extended time decreased missed cleavage rates.

**Table S1.** Optimizations of gel-making conditions.

| **Gel No.** | **Anchoring time** | **Hydrogel composition**  **(SMA:PAE)** | **SDS concentration in homogenization buffer** | **Homogenization condition** |
| --- | --- | --- | --- | --- |
| Gel 1 | 1 h | 71:1 | 20% | 105°C, 1 h  (0.02-0.025 MPa) |
| Gel 2 | 1 h | 156:1 | 20% |  |
| Gel 3 | 1 h | 71:1 | 5.8% | 110°C, 1 h  (0.045 MPa) |
| Gel 4 | 1 h | 71:1 | 10% |  |

**Table S2. Optimizations of reduction and alkylation steps as well as in-gel proteolytic digestion.**

| **Gel No.** | **Reduction and alkylation steps** | **Enzyme digestion** |
| --- | --- | --- |
| 1 | Before embedding (whole gel) | Trypsin 12.5 ng/µL (4 h) +  Trypsin 12.5 ng/µL (overnight) |
| 2 | After homogenization (whole gel) |  |
| 3 | During in-gel digestion (single punch) |  |
|  | During in-gel digestion (single punch) | Lys-C 6.25 ng/µL (4 h) +  Trypsin 12.5 ng/µL (overnight) |
|  | During in-gel digestion (single punch) | Trypsin 12.5 ng/µL (3 h) |

Note that these optimizations are carried using mouse brain slices under the ProteomEx workflow.

**Table S3.** MS settings for DIA-PASEF.

| **ExpType** | **Repetitions** | **KA** | **m1** | **m2** | **CEA** | **KB** | **m3** | **m4** | **CEB** | **Steps** |
| --- | --- | --- | --- | --- | --- | --- | --- | --- | --- | --- |
| MS1 | 1 | - | - | - | - | - | - | - | - | - |
| PASEF | 1 | 0.703 | 384 | 412 | -1 | 1.119 | 909 | -1 | -1 | 4 |
| PASEF | 1 | 0.717 | 409 | 437 | -1 | 1.133 | 934 | -1 | -1 | 4 |
| PASEF | 1 | 0.731 | 434 | 462 | -1 | 1.147 | 959 | -1 | -1 | 4 |
| PASEF | 1 | 0.745 | 459 | 487 | -1 | 1.161 | 984 | -1 | -1 | 4 |
| PASEF | 1 | 0.759 | 484 | 512 | -1 | 1.175 | 1009 | -1 | -1 | 4 |
| PASEF | 1 | 0.774 | 509 | 537 | -1 | 1.189 | 1034 | -1 | -1 | 4 |
| PASEF | 1 | 0.788 | 534 | 562 | -1 | 1.203 | 1059 | -1 | -1 | 4 |
| PASEF | 1 | 0.797 | 384 | 412 | -1 | 1.213 | 909 | -1 | -1 | 4 |
| PASEF | 1 | 0.811 | 409 | 437 | -1 | 1.227 | 934 | -1 | -1 | 4 |
| PASEF | 1 | 0.825 | 434 | 462 | -1 | 1.241 | 959 | -1 | -1 | 4 |
| PASEF | 1 | 0.839 | 459 | 487 | -1 | 1.255 | 984 | -1 | -1 | 4 |
| PASEF | 1 | 0.853 | 484 | 512 | -1 | 1.269 | 1009 | -1 | -1 | 4 |
| PASEF | 1 | 0.867 | 509 | 537 | -1 | 1.283 | 1034 | -1 | -1 | 4 |
| PASEF | 1 | 0.881 | 534 | 562 | -1 | 1.297 | 1059 | -1 | -1 | 4 |

**Table S4.** Chemical composition of reagents and buffers for FAXP.

| **Purpose** | **Solution name** | **Solution formula** | **Storage temperature** |
| --- | --- | --- | --- |
| Protein Anchoring | NSA stock | Dissolve 100 mg of NSA in the anhydrous DMSO to make the final volume at 10 mL. | Store at 4℃ |
|  | Protein anchoring buffer | Mix 25 mL 0.2 M MES stock solution with 25 mL ddH_2_O to make the final volume at 50 mL and test if the pH is at 6.0. | Store at 4℃ |
|  | Protein anchoring solution | 2 μL NSA stock solution with 198 μL protein anchoring buffer. | Freshly prepared |
|  | Anchoring termination buffer | Mix 2 mL 0.5 M MOPS stock solution with 8 mL ddH_2_O to make the final volume at 10 mL and test if the pH is at 7.0. | Store at 4℃ |
| Gelation | PAE stock solution | Dissolve 1.15 g PAE in the tetrahydrofuran (THF) to make the final volume at 10 mL and avoid light. | Store at 4℃ |
|  | Monomer solution stock | 10 mL, pH 6.5;  3.137 g N,N-dimethylacrylamide  (DMAA) (~3 mL);  0.8624 g Sodium methacrylate (SMA);  269.5 μL Pentaerythritol allyl ether (PAE stock solution);  ddH2O 4.380 mL;  10% HCl 350 μL | Store at 4℃ |
|  | APS stock | Dissolve 0.1 g APS in ddH_2_O to make the final volume at 1 mL. | Freshly prepared |
|  | TEMED stock | Dissolve 0.1 g TEMED (about 0.129 mL) in ddH_2_O (0.871) to make the final volume at 1 mL. | Store at 4℃ |
|  | Activated monomer solution | 1 mL - pH 6.5;  900 μL Monomer solution stock;  30 μL APS stock;  20 μL TEMED stock;  50 μL ddH_2_O | Freshly prepared |
| Homogenization | Protein denaturation buffer | 200 mL - pH 7.0;  10 mL Tris, 1 M, pH 8;  40 g, SDS;  1.8612 g, EDTA-Na_2_·2H_2_O;  8 mL, NaCl, 5 M | Store at RT |
| Reduction and alkylation | 20 mM TCEP solution | Dissolve 0.1433 g TCEP in 1× PBS (pH 7.4) to make the final volume at 25 mL. | Freshly prepared |
|  | 45 mM TCEP solution | Dissolve 0.3225 g TCEP in 1× PBS (pH 7.4) to make the final volume at 25 mL. | Freshly prepared |
|  | 10 mM DTT solution | Dissolve 0.03856 g DTT in 1× PBS (pH 7.4) to make the final volume at 25 mL. | Freshly prepared |
|  | 20 mM DTT solution | Dissolve 0.07712 g DTT in 1× PBS (pH 7.4) to make the final volume at 25 mL. | Freshly prepared |
|  | 55 mM IAA solution | Dissolve 0.25432 g IAA in 1× PBS (pH 7.4) to make the final volume at 25 mL. | Freshly prepared |
|  | 100 mM IAA solution | Dissolve 0.4624 g IAA in 1× PBS (pH 7.4) to make the final volume at 25 mL. | Freshly prepared |
| Coomassie blue staining | Coomassie Blue solution | Mix R-250 with ddH_2_O to a final density of 1 mg/mL. | Store at RT |
| Enzymatic digestion | 80% ACN | Mix 8 mL ACN with 2 mL ddH_2_O to make the final volume at 10 mL. | Store at RT |
|  | 100 mM ABB | Dissolve 79.06 mg ABB in the ddH_2_O to make the final volume at 10 mL. | Store at RT |
|  | 50 mM ABB | Mix 1 mL of 100 mM ABB with 1 mL of ddH_2_O to make final volume at 2 mL. | Store at RT |
|  | 50% ACN/50% 100 mM ABB | Mix 10 mL of 100 mM ABB with 10 mL of ACN to make final volume at 20 mL. | Store at RT |
|  | 50% ACN/50% ddH_2_O | Mix 10 mL of ACN with 10 mL of ddH_2_O to make final volume at 20 mL. | Store at RT |
|  | Trypsin stock solution | Add 200 μL of 0.1% HAc (pH = 3) to 100 μg trypsin, aliquot into 50 μL/vial. | Store at -20℃ |
|  | Digestion buffer 1 | Mix 0.5 mL of ACN with 4.5 mL of 50 mM ABB to make the final volume at 5 mL. | Freshly prepared |
|  | Digestion buffer 2 | Mix 0.1 mL of 100 mM ABB with 0.9 mL of ddH_2_O to make final the volume at 1 mL. | Freshly prepared |
|  | Working solution for in-tip digestion | Add 5 μL trypsin stock solution to 195 μL of digestion buffer 1. Need to test the pH by stripes, which should be around 8-9. | Freshly prepared |
|  | Working solution for in-gel digestion | Add 5 μL trypsin stock solution to 195 μL of digestion buffer 1. Need to test the pH by stripes, which should be around 8-9. | Freshly prepared |
| Peptide extraction | Solvent buffer A – in-tip digestion | Mix 2 μL ACN with 10 μL Trifluoroacetic acid (TFA) and 9.8 mL ddH_2_O to make the final volume at 10 mL. | Store at RT |
|  | Solvent buffer B – in-tip digestion | Mix 7 mL ACN with 10 μL TFA and 3 mL ddH_2_O to make the final volume at 10 mL. | Store at RT |
|  | Solvent buffer A – in-gel digestion | 25 mM ABB solution; | Store at RT |
|  | Solvent buffer B – in-gel digestion | 40 mL;  20 mL ACN;  1 mL formic acid (FA);  Add 19 mL ddH_2_O to 40 mL; | Store at RT |
|  | Solvent buffer C – in-gel digestion | 100% ACN; | Store at RT |
| Desalting | Activation buffer | 100% Methanol; | Store at RT |
|  | Equilibration buffer A | 10 mL;  8 mL ACN;  10 μL TFA;  Add ddH_2_O to 10 mL | Store at RT |
|  | Equilibration buffer/washing buffer | 10 mL;  0.2 mL ACN;  10 μL TFA;  Add ddH_2_O to 10 mL; | Store at RT |
|  | Elution buffer | 10 mL;  4 mL ACN;  10 μL TFA;  Add ddH_2_O to 10 mL; | Store at RT |
| Loading for MS acquisition | MS buffer | 10 mL;  0.2 mL ACN;  10 μL FA;  Add ddH_2_O to 10 mL; | Store at 4℃ |

**Table S5.** Main chemicals, reagents, consumables, and instruments used for FAXP and their corresponding suppliers and catalog numbers.

|  | **Name** | **Supplier** | **Catalog number** |
| --- | --- | --- | --- |
| *Reagents* | | | |
| H&E staining | Hematoxylin and eosin staining solution | Shanghai Yuanye | R20570-2*100ml |
|  | Differentiation solution (acid alcohol; 1%) | Shanghai Yuanye | R20778-500ml |
|  | Scott's solution | Shanghai Yuanye | R20596-500ml |
|  | Neutral balsam | Macklin | N861409-25ml |
| Protein anchoring | N-succinimidyl acrylate (NSA) | TCI | S0814 |
|  | DMSO | Sigma-Aldrich | D5879-100ML |
|  | MOPS, 0.5M, pH 7.0 | Macklin | M885700 |
|  | MES, 0.2M, pH 6.0 | Macklin | M885671 |
| Gelation | Pentaerythritol allyl ether (PAE) | Sigma-Aldrich | 251720-100G |
|  | Tetrahydrofuran (THF), HPLC | Macklin | T818769-500ml |
|  | N, N-dimethylacrylamide (DMAA) | Sigma-Aldrich | 274135-500ML |
|  | Sodium methacrylate (SMA) | Sigma-Aldrich | 408212-50G |
|  | Ammonium persulfate (APS) | Sigma-Aldrich | A3678 |
|  | N, N,N′, N′-Tetramethylethylenediamine (TEMED, density 0.775 g/mL) | Sigma-Aldrich | T7024 |
| Homogenization | Sodium dodecyl sulfate (SDS) | Macklin | S817790-500g |
|  | EDTA-Na_2_·2H_2_O | Sigma-Aldrich | E4884-100g |
|  | Tris base | Sigma-Aldrich | T6791-500G |
|  | NaCl | Sigma-Aldrich | S9888-500G |
| Reduction and alkylation | Tris(2-carboxyethyl)phosphine (TCEP) | Adamas Reagent | 61820E |
|  | Dithiothreitol (DTT) | Thermo Fisher | R0861 |
|  | Iodoacetamide (IAA) | Sigma-Aldrich | I6125-25G |
| Staining | Brilliant blue R-250 | BBI | A610037-025 |
|  | SYPRO Ruby | Thermo Fisher | S12000 |
| Enzymatic digestion and peptide elution | Formic acid (FA) | Thermo Fisher | A117-50 |
|  | Methanol | General-reagent | G75851B |
|  | Lys-C | Hualishi | HLS LYS001C-50μg |
|  | Trypsin | Hualishi | HLS TRY001C-100μg |
|  | Ammonium bicarbonate (ABB) | Sigma-Aldrich | A6141-1kg |
|  | Acetonitrile (ACN) | General-reagent | G80988B |
|  |  | Thermo Fisher | A955-4 |
|  | MS grade water | Thermo Fisher | W6-4 |
|  | Trifluoroacetic acid (TFA) | Adamas Reagent | 81548K |
| *Consumables* | | | |
|  | 200 μm in thickness × 1.5 mm in diameter glass chamber | Customized |  |
|  | Single Specimen \| No. 1.5 Coverslip \| 98mm X 67mm Viewing Area \| Uncoated | MatTek | P384G-1.5-10872-C |
|  | 6 cm diameter petri dish | NEST | 705001 |
|  | 9 cm diameter petri dish | NEST | 752004 |
|  | 1.5 ml EP tube | Axygen | MCT-150-C |
|  | Empore C18 47mm Extraction Disks | 3M | 66883-U |
|  | Low protein binding tubes, 1.5 mL | Thermo Fisher | 90410 |
|  | QSP standard tips | Thermo Fisher | TLR102-Q |
| *Instruments* | | | |
|  | Fluorescence Stereo Zoom Microscope | ZEISS | Axio Zoom.V16 |
|  | Vacuum drying oven | Shanghai YiHeng | DZF-6032 |
|  | Autoclave | TOMY | SX-700 |
